# 3D, multi-omic imaging reveals molecular biomarkers of the pre-metastatic niche in lung cancer

**DOI:** 10.64898/2026.02.18.706515

**Authors:** John Michel, André Forjaz, Vasco Queiroga, Kelly Casella, Kaitlin Stivers, Helen Nguyen, Maria Browne, Feiyu Chen, Ada Tam, Omkar Dhaygude, Hongni Fan, Hitoaki Maehira, Cheng Ting Lin, Elise Gray-Gaillard, Talia L. Benducci, Suguru Yamauchi, Ife Shoyombo, Ziying Xu, Eugene Shenderov, Peng Huang, Denis Wirtz, Jennifer H. Elisseeff, Yun Chen, Malcolm V. Brock, Ashley L. Kiemen, Franck Housseau

## Abstract

The recurrence rate following complete surgical resection of primary non-small cell lung cancer is as high as 55%, yet no approach currently exists to evaluate the risk of local recurrence. The premetastatic paradigm is the recognition that metastasis is preceded by reprogramming naïve tissues to prime a microenvironment for tumor cell survival and subsequent reactivation. Identification of biomarkers of the pre-metastatic niche would allow us to evaluate a patient’s risk of local relapse in the normal lung parenchyma surrounding the resected tumor. We designed a workflow incorporating *in vivo* modelling, radiology, and deep learning-guided three-dimensional (3D) imaging, spatial proteomics, and transcriptomics to identify previously unreported signals associated with the early transformation of the lung parenchyma announcing regional metastasis. We curated biorepository spanning timepoints before and after resection of primary Lewis Lung Carcinoma (LLC) tumors. Using radiology and cellular resolution 3D histology, we calculated the number and distribution of metastases in mouse lungs and developed an algorithm to guide placement of spatial proteomics and transcriptomics to regions containing early micro-metastases and the pre-metastatic microenvironment. Molecular and tissue features associated with presence, size, and location of metastases guided the identification of both myeloid (F4/80) and senescent (p16/p21) cell signatures in the premetastatic and metastatic environments. Finally, multiparametric flow cytometry of metastatic lungs in a senescence reporter GEMM (tdTomato-p16 INKA mice) resolved senescent cells including alveolar macrophages as the cellular phenotypes associated with these early premetastatic signatures. Altogether, this work highlights a novel AI-assisted approach for detection of biomarkers of tissue remodeling during lung cancer invasion.

## Introduction

Non-small cell lung cancer (NSCLC) is the most common form of lung cancer. Metastasis is a major driver of mortality and morbidity in NSCLC following surgical resection of a primary tumor with curative intent.^1^ Pulmonary metastases occur with an incidence of ∼30% in those with malignant cancer; further, these patients have a ∼35% 5-year survival rate once metastasis occurs.^2–4^ Therefore, our ability to effectively treat lung cancer is limited by our inability to identify metastases early and inhibit regional metastasis, emphasizing the need for further work to clarify the early premetastatic mechanisms in distant organs which are responsible for further cancer dissemination.

The paradigm of the premetastatic niche (PreMN) proposes that certain organs are selectively primed for metastasis through discrete tissue remodeling preceding the arrival of tumor cells.^5^ Many markers of metastatic cascade, such as tumor secreted factors, bone marrow derived myeloid cell recruitment, and tumor derived exosomes, have been identified among others features at the bulk level of a whole organ by general screening for expression compared to a non-metastatic environment, without however delineating PreMN and limiting therefore their clinical utility in predicting metastatic site and risk.^6^ Novel multi-omics (transcriptomic, proteomic, imaging) approaches allow the resolution of the PreMN as spatially infrequent tissue alterations that proceed metastatic foci formation.^7^ To overcome the challenges inherent to identifying rare and subtle tissue alterations of the Pre-MN, we integrated novel, artificial intelligence (AI)-guided image analysis tools with spatial transcriptomics and multiplexed IF.^8^ A better understanding of premetastatic events in the lung parenchyma distant from resectable primary tumors in NSCLC patients may improve our means to detect and intercept metastases formation via reversing early microenvironmental priming as we previously showed.^9^

We optimized a novel pipeline combining *in vivo* modelling and whole organ, multi-omic imaging to leverage a previously established preclinical model of pulmonary metastases formation post Lewis Lung Carcinoma (LLC) surgical resection in profiling spatially and molecularly the earliest stromal signs of metastasis cultivation in the lungs. To isolate the effects of the circulating tumor cells and micro-metastases, we surgically resected the primary tumors and collected whole murine lungs at different timepoints in the days surrounding primary tumor resection. As micro-metastases are sparsely distributed within the lung parenchyma, a three-dimensional (3D) approach is essential to capture their spatial context and identify rare, microscopic early metastatic colonization.^10^ We used CODA, an AI-based platform for cellular resolution, 3D tissue reconstruction to generate quantitative anatomical maps of the pulmonary anatomy.^11–14^ With these 3D maps, we resolved the number, volume, and location of individual metastases across the lung, identifying micro-metastases that were not visible in pre-surgical computed tomography (CT) imaging. We strategically applied spatial proteomic and transcriptomic profiling at regions of interest containing micro-metastases to reveal distinct populations of senescent alveolar macrophages and stromal reorganization as preferentially increased around micro-metastasis. This knowledge permitted us to ask specific questions regarding how metastases with given sets of features influenced their environment, which is not possible with the traditional methods. For example, how tumor volume and distance from a tumor impact the transcriptomic profile of adjacent lung tissue.

In summary, we successfully developed a platform to probe whole organs to identify rare regions of early micro-metastasis. With this platform we identified novel candidate biomarkers of lung parenchyma reorganization, which may improve our ability to detect PreMN and predict recurrence risk in NSCLC patients.

## Results

### Myeloid cells are enriched in the metastatic lungs of a syngeneic mouse model of metastatic lung cancer

To discern changes in the immune cell populations associated with the metastatic cascade, we designed an *in vivo* workflow to harvest murine lungs perturbed by the presence of primary lung tumors. We identified the Lewis lung carcinoma (LLC) only-in-mice model as a well-validated tumor implantation and resection model of metastasis formation.^15,16^ By incorporating the effect of surgical resection, and therefore the systemic response to wound healing, the LLC model recapitulates the clinically and physiologically relevant impact of the injury that is generated when a tumor is removed from a patient, and the subsequent metastatic burst. (**Fig. 1a**) We harvested lungs seven days following tumor resection, a time point where metastases are detectable using conventional CT-scans. We ran high parameter flow cytometry to identify immune cell types differentially represented in age and sex-matched mouse lungs that underwent tumor implantation and resection surgery (Tumor group); lungs that underwent identical surgery with no tumor (Mock group); and control lungs from mice that did not undergo surgery or tumor exposure (Naïve group). Lungs were harvested from the three groups processed, and a comparative comprehensive flow cytometry analysis was performed to characterize the immune cell populations. (**Fig. S1a**) Both a pan-immune and intracellular cytokine panel were run on the resulting cells to identify shifts in cell types, abundance, and functional status.

**Fig. 1.**
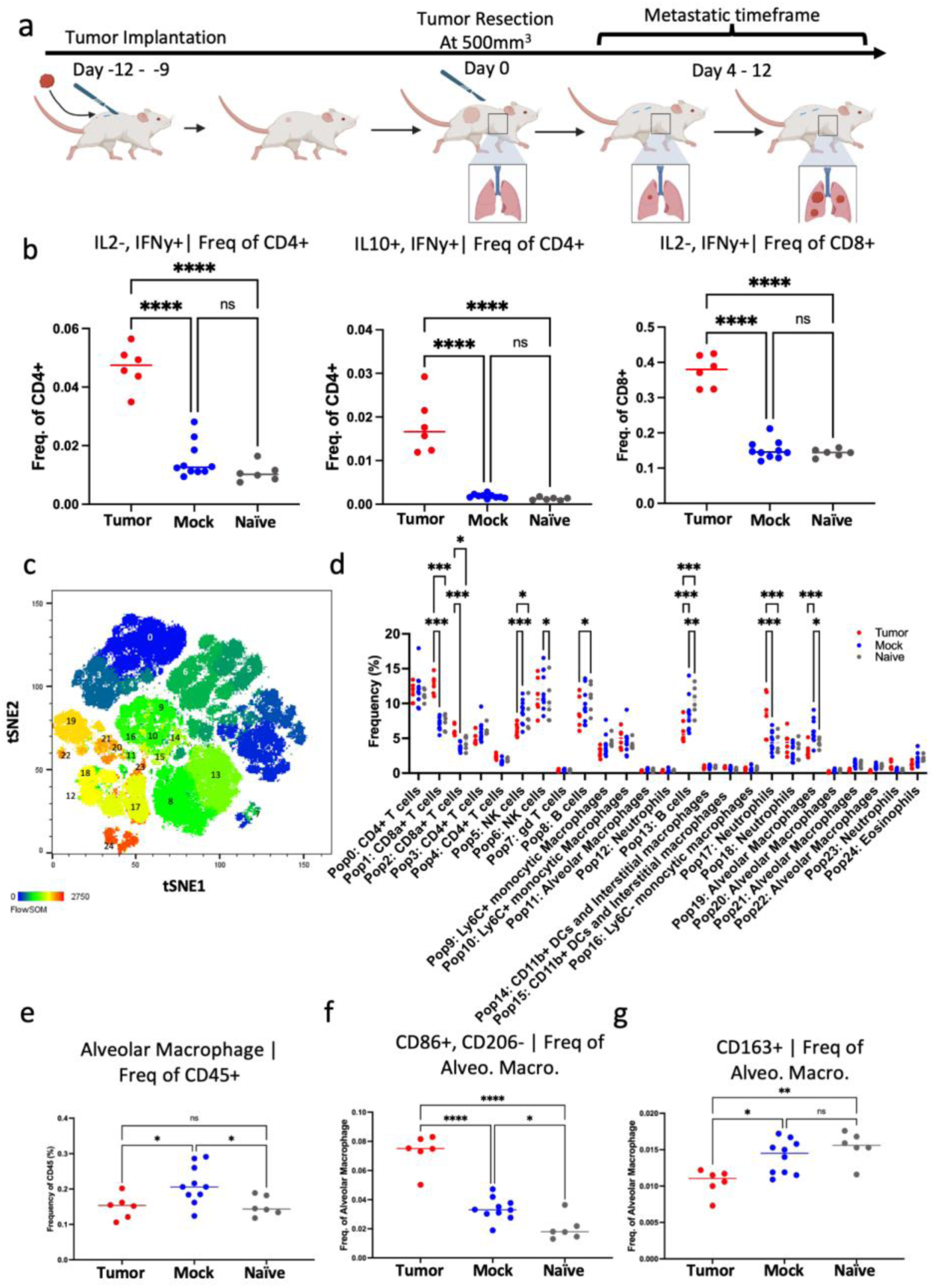
High parameter flow analysis reveals skewing of the lymphoid and myeloid compartments. **(a)** Description of typical timeline for LLC tumor implantation experiment **(b)** Flow comparison of IL2^-^ IFNy^+^ CD4^+^ T cells, IL10^+^ IFNy^+^ CD4^+^ T cells, and IL2^-^ IFNy^+^ CD8^+^ T cells as frequency of parent population (n = 6 tumor; n = 10 mock; n = 6 naïve) **(c)** tSNE representation of entire pan-immune flow data **(d)** Comparison of tSNE representations between tumor, mock, and naïve groups **(e)** Flow comparison of alveolar macrophage, CD86^+^ CD206^-^ alveolar macrophages, and CD163^+^ alveolar macrophages as frequency of parent population (n = 6 tumor; n = 10 mock; n = 6 naïve) (**Statistics**) No statistical analysis performed for pan-immune tSNE plots, or experimental outline (a, c) Flow cytometry comparisons: ordinary one-way ANOVA with Tukey’s multiple comparisons test (only relevant comparisons shown) (b, e) Two-way ANOVA with Tukey’s multiple comparisons test (only relevant comparisons shown) (d) * p<0.05, ** p<0.01, *** p<0.001, **** p<0.0001.

In the Tumor group, CD4^+^ and CD8^+^ T cell populations had significant changes in cytokine expression. (**Fig. 1b, S2a**) Both control groups (Mock and Naïve mice) exhibited similar proportions of IFNy and IL10 producing CD4^+^ cells. A higher proportion of CD4^+^ cells produced IFNy and IL10 in the Tumor condition compared to both control groups. Further, an increased percentage of CD8^+^ T cells in Tumor condition produced IFNy compared with the two controls. These results validated the impact of metastasis on the local immune environment with a predominant type-1 inflammatory environment in the tumor bearing mice as compared to the control mice.^17^

Next, an unsupervised analysis of CD45^+^ immune cell types in each group (Tumor, Mock, Naïve) was performed using the tSNE and flowSOM (FlowJo) platform to generate 25 clusters. (**Fig.1c, 1d, S1b**) In accordance with the flow cytometry analysis, we identified an increase in CD8+ T cells in the Tumor condition (Pop 1, 2 in **Fig.1d**). In the myeloid compartment, a neutrophil population was increased in the Tumor condition as compared to Mock and Naive (Pop17 in **Fig.1d**). These finding corroborated our previous observation in this LLC model where PreMN formation was associated with increased infiltrating myeloid derived suppressor cells (MDSCs) before the detection of large metastases in the lungs.^15^ Further, an alveolar macrophage population (Pop 19 in **Fig.1d**; feature plots of key markers in **Fig. S1c**) was significantly increased in the Mock condition as compared to Tumor or Naïve ones. Exploration of the alteration of the myeloid landscape showed a striking difference in alveolar macrophages (CD45^+^ CD11b^mid-low^ SigF^+^ CD11c^+^ CD64^+^) frequency between both groups. (**Fig. 1e, S2b**). Although no change was observed in alveolar macrophage abundance, we observed an increase of a CD86^+^, CD206^-^ subset along with a decrease of CD163^+^ alveolar macrophages in the tumor condition, suggesting a functional macrophage rewiring in the premetastatic environment. (**Fig. 1f, 1g)** There were minimal differences between the Mock and Naïve conditions, dissociating tumor intrinsic from systemic wound healing signals in orchestrating alveolar macrophage. tSNE alveolar macrophage visualization confirmed a phenotypic shift in the Tumor condition as compared to Naive and Mock – towards CD86+ expression in the Tumor condition. (**Fig. S1d, S1e**)

### AI-assisted 3D pathology imaging outperforms conventional thoracic CT in detection of pulmonary micro-metastases

Following characterization of immune evolution in the LLC model, we generated a biorepository of lungs spanning the pre-metastatic and metastatic stages to enable capture of the changes to the structural, cellular, and extra-cellular matrix (ECM). We harvested lungs three days before LLC tumor resection D-3 (n=2) as well as D2 (n=4), D4 (n=4), D7(n=4), and D11 (n=2) after tumor resection. The effect of surgery only was controlled using Mock group as described earlier. Each mouse was first monitored for metastasis using conventional thoracic CT scans and analysis by a trained radiologist. We detected tumors on CT scans as early as day four post primary tumor resection and most of the mice were positive by day seven (**Fig.2**).

**Fig. 2.**
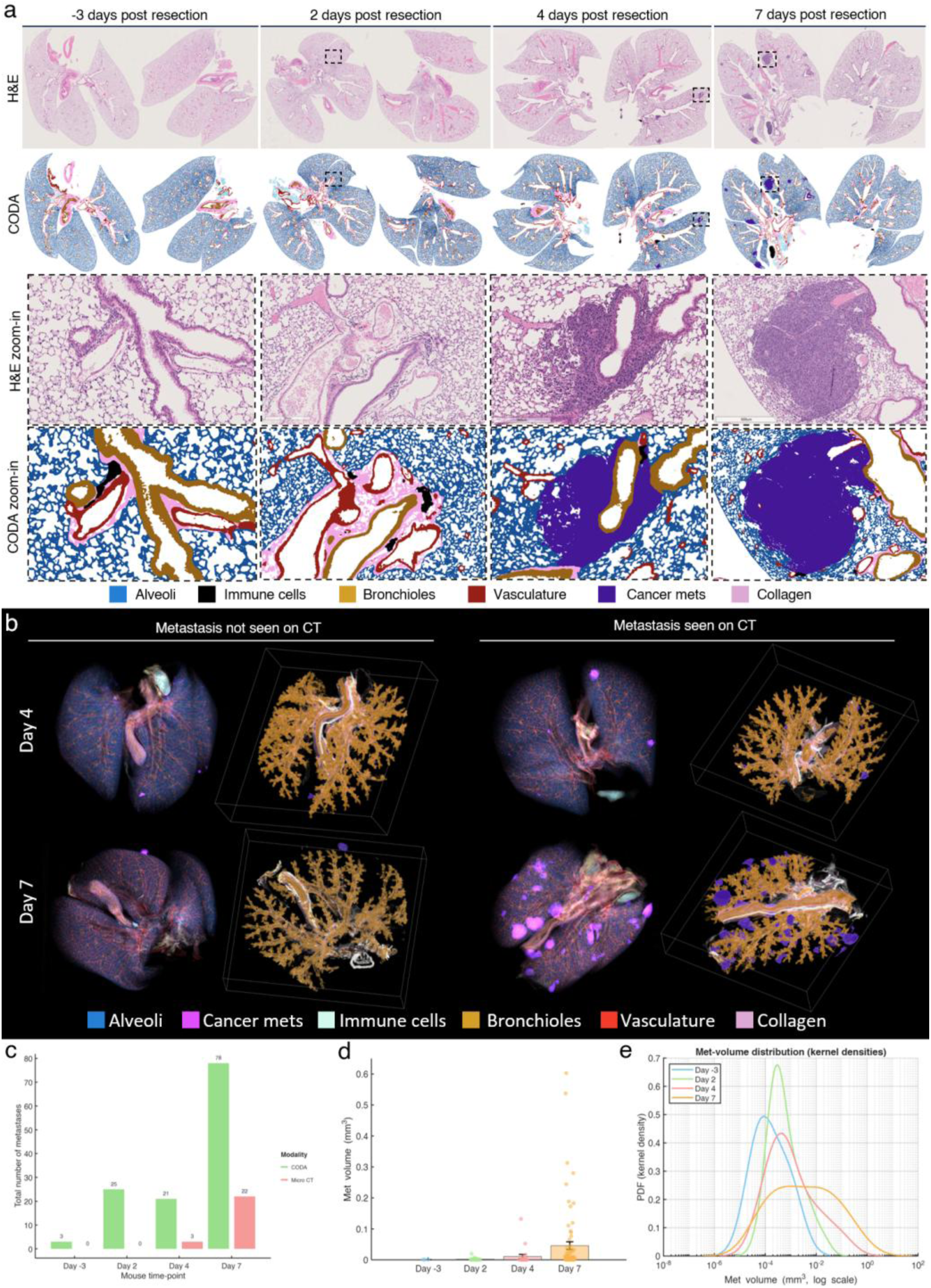
Mapping lung microanatomy and metastasis using 3D CODA-guided pathology. **(a)** Serial sectioning and H&E-staining of 300 sections per mouse lung at days -3, 2, 4 and 7 post-resection. Deep learning-based segmentation of microanatomical tissues in mouse lungs. **(b)** CT-scans were collected for each mouse lung resection. Expert-screened CT-scans missed microscopic metastases that were captured by 3D-CODA single cell mapping. **(c)** Number of metastases detected in CT and using CODA for each mouse at different days post-resection (n = 2 mice per timepoint). **(d)** Metastasis volumes detected using CODA at different days post-resection (n = 2 mice per timepoint). **(e)** Comparison of metastasis volume distributions for each day. Kernel density estimation reveals mostly smaller metastases across early timepoints (day - 3 and day 2), and larger metastases in later time points (day4 and day 7).

The lungs were then collected, formalin fixed following controlled inflation, paraffin embedded, serially sectioned, histologically H&E stained, and scanned at high-resolution for 3D reconstructed using the AI-based platform CODA (**Fig.2a, S3a**).^11^ Whole lung reconstruction at single cell resolution level was performed on blocks without detectable metastases by radiomics, for timepoints D-3 (n=2), 2 (n=2), 4 (n=2), and 7 (n=2). (**Fig. 2a, 2b**) CODA identified 9 tissue types, consisting of cancer cells, bronchioles, alveoli, blood vessels, bronchiole, stroma, cartilage, inflammation, and nerves, with an overall model accuracy of 97.3% **(Fig. S3b)**

Since LLC cells are Vimentin^+^ CK19^-^ automated detection of cancer cells using the CODA segmentation algorithm was validated using IHC for Vimentin (EMT+ LLC cells) versus CD45/CD31/CK19 staining performed on adjacent slides **(Fig. S3a)**.^18^ Regions with metastasis indexed by CODA were characterized by the absence of CD45, CD31, and CK19 staining, and by the presence of a strong Vimentin signal. **(Fig. S3c and d)**. Our results show the ability of the CODA segmentation model to accurately segment metastatic lesions as small as 3.63x10^4^ µm³.

Using CODA, we detected metastasis in all mice across all timepoints, even when conventional CT analysis of the mice did not detect them at earlier timepoints (days -3 and 4), confirming the high sensitivity of AI-based 3D pathology imaging (**Fig. 2c).** Further, reconstruction of these tissue features in 3D demonstrated preservation of the underlying morphology. In total, CT scan imaging identified a total of 25 metastases and CODA identified 127 metastases in all eight mice (**Fig. 2c)**. The distribution of detected metastases varied by day, with 3 metastases recorded on day -3, 25 on day 2, 21 on day 4, and a significant increase to 78 on day 7 (n=2 per time point). Mean and median metastatic volumes increased across time points, highlighting also the sensitivity of CODA in detection of early metastases.^18^ **(Fig. 2d-e; Sup. Table 1)**

### CODA-guided spatial transcriptomics and proteomics to define molecular and histological features of premetastatic lungs

The 3D characterization of metastasis-bearing lungs at single cell resolution level being now available, we designed a workflow to pinpoint regions of interest (ROIs) in whole lung tissues likely to contain the molecular and histological biomarkers characterizing early metastatic pulmonary tissues, where micro-metastases are not detectable by conventional thoracic CT-scans. Three histology features were used to define ROIs for targeted multi-omics: 1) presence; 2) proximity; and 3) size of metastases. Our rational was that microenvironments at the margin of the micro-metastases had a high probability to be enriched in signals associated with pulmonary parenchyma reorganization prior to disseminated tumor cell settlement.

We chose a D4 post-resection sample, which constituted the earliest timepoint at which we found both micro- and macro-metastases. Of the available slides, we narrowed our query down to the image in which micro-metastasis area was maximized, as defined using the 3D AI-labelled imaging (**Fig. 3a, left**). Within the target image, we calculated the 3D Euclidean distance to the nearest micro-or macro metastasis, as well as the 3D Euclidian distance to the nearest blood vessel and represented this data topographically. We then defined 24 ROIs per slide according to their distance from metastases as tumor (*d*≤0.5mm), near (0.5mm>*d* ≤1.5mm) and far (1.5mm>*d*≤2.5mm) (**Fig. 3a, right**).

**Fig. 3.**
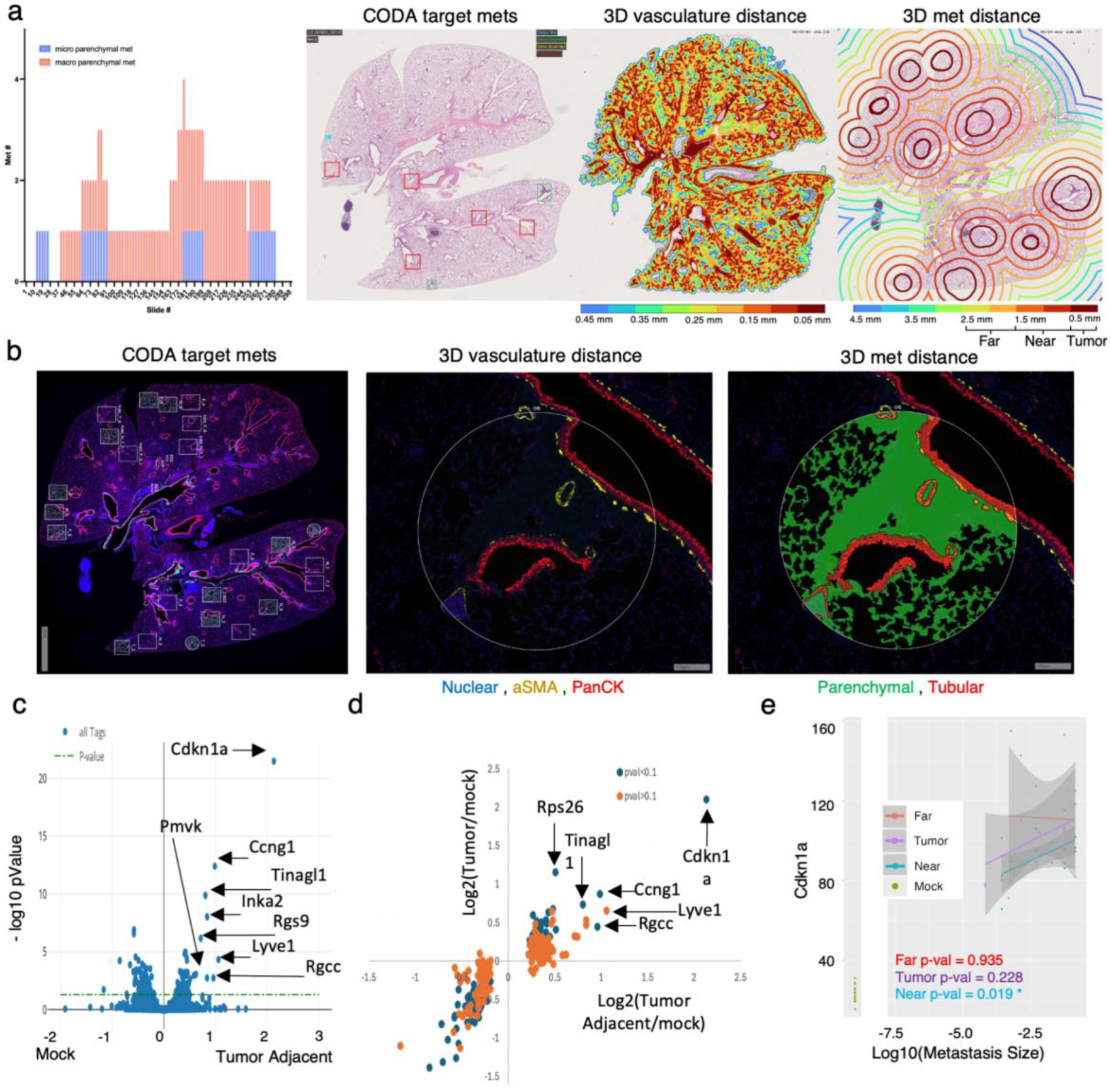
GeoMx analysis of the metastatic environment based on CODA, ROI Identification. **(a)** (right) Number of micro and macro-metastases identified per slide by CODA and representative slide with both micro and macro metastases (blue = micro; green = macro). (left) Overlay of topographical map of 3D distance from nearest blood vessel. Overlay of topographical map of 3D distance from nearest metastasis **(b)** Representative example of metastasis on GeoMx imaging software and segment annotation based on Nuclear, aSMA, and PanCK staining **(c)** Volcano plot (-log10 pValue vs Log2) comparison of tumor adjacent vs Mock annotation for all genes. Significant genes (p<0.05) with high log2 fold change were annotated. **(d)** Scatter plot of genes with padj<0.1 from the tumor adjacent vs mock annotation comparison, plotted based on tumor vs mock annotation comparison. X-axis: log2(tumor adjacent/mock) and y-axis: log2(tumor/ mock). Blue indicated padj<0.1 in tumor vs mock annotation comparison. Orange indicates padj>0.1 in tumor vs mock annotation comparison. **(e)** Scatter plot representing Cdkn1a expression (count) per GeoMx Parenchymal ROI, with Pearson correlation shown. (**Statistics**) No statistical analysis was performed tumor quantification graphs or histological images (a,b) Log2 and padj values were generated using a linear mixed model: ∼ Tags + (1+ Tags | Scan_ID) + (1+ Tags | Scan_ID : Mouse) with Benjamini-Hochberg correction. Tags corresponds to each sample, Scan_ID to the slide from which the data was collected, and mouse to the mouse which produced the slides. For near vs far comparison, near vs far annotations were separated in the Tags of the model. For all other analyses these annotations were combined into a tumor adjacent annotation in the Tags of the model (c).

ROIs from a timepoint-matched mock surgery mouse were included as reference. GeoMx spatial digital profiler NanoString platform delineated ROIs and their cell type content by IF (area of interest or AOI). Each AOI can be then selectively analyzed by transcriptomics. We discriminated 2 AOI per ROI, parenchymal (alveolar regions Syto83^+^ PanCK^-^ aSMA^-^), acquired first, and tubular (SMA^+^ or PanCK^+^) regions. Tubular structures correspond to large branches of the vasculature and airway structures larger than a terminal bronchiole, arteriole, or venule.^19^ (**Fig. 3b**). To minimize sample bias and facilitate the standardization of our analysis, we excluded these tubular structures due to their sporadic and spare distribution in the lungs. We, therefore, performed our transcriptomic analysis on the parenchymal segments, which had a higher number of samples and a greater overall cell count.

Our results first highlighted a statistically significant impact of the metastasis size (micro-versus macro-metastases, as defined using a 1.5 mm^3^ volume threshold, see materials and methods) on GSEA pathway using the Reactome genesets and including *immune system function* such as *Neutrophil Degranulation* in large tumors. **(Fig. S4a)** This result matches our flow cytometry data **(Fig.1d)**, which showed the accumulation of myeloid populations (in particular neutrophils) in tumor bearing mice at later time points. Leading edge of the signature genes contained defensins (including Defa34) and serine protease inhibitors (including Serpinb1a), which may act in this context as an anti-apoptotic signal. *Siglecf*, a key marker of alveolar macrophages, was also detected.

A similar analysis between ROIs Near and Far from a metastasis (as calculated in 3D space) pointed next towards an enrichment of immune system function and extracellular matrix organization signature in the far ROIs. **(Fig. S4b)** Interestingly neutrophil degranulation signature was the most upregulated of the immune system function genesets and leading-edge genes of the ECM geneset includes collagens (Col8a2), Loxl4 (lung fibrosis), and Scube3 (EMT). This result suggested a potential active remodeling in the Far regions, as opposed to Near regions, which already have been reorganized. Transposed to premetastatic microenvironment concept, these Far ROI-enriched signatures may be biomarker candidates for PreMN formation.

We next sought to determine whether the presence of a metastatic lesion impacts biomarkers identified in the lung stroma. We combined the *Near* and *Far* annotations into a single category, which we termed Tumor adjacent. Differential gene expression analysis between the parenchymal segments in Tumor adjacent versus tissue-matched ROIs in Mock pulmonary tissue identified multiple significantly upregulated markers in the tumor adjacent annotation, including *Ccng1* and *Cdkn1a* coding p21, markers of cellular senescence *Tinagl1*, and *Lyve1*, markers of angiogenesis, or *Rgcc* coding RGC-32 associated with pulmonary fibrosis. (**Fig. 3c)** ^20,21^ These markers indicate that tissue reorganization is highly relevant to PreMN formation. Furthermore, Lyve1 is a marker of angiogenesis expressed by macrophages involved in vasculature fitness. Senescent alveolar macrophages have previously been associated with the development of primary tumors in the lung environment.^22,23^ Interestingly, when comparing tumor and tumor adjacent to the Mock tissues for these marker’s expression, there was no statistical significance (p=0.1) in differential expression for the expression of Lyve1, but *Cdkn1a* remained statistically differentially expressed (p<0.05) between Tum and Tum adjacent. (**Fig. 3d)** Finally, when we once again considered the role of size of metastasis on expression of *Cdkn1a*, we noted that tumors with larger size had higher expression of *Cdkn1a* in Near regions. (**Fig. 3e)**

### Senescent cells are associated with the premetastatic micro-environment

Based on the results of the GeoMx analysis we aimed to confirm the detection of cellular senescence markers in the LLC metastatic environment. Both p16 and p21 are classical markers of senescence and can be expressed by nearly any cell type. To rule out senescent DTC, we implanted LLC tumors in the p16-CreERT2;Ai14 (INKA) mouse model.^24^ In this model, only p16^+^ cells from the INKA mice express tdTomato following exposure to Tamoxifen. We harvested and processed lung tissues according to the kinetic established earlier and ran flow cytometry to compare Tamoxifen (p16+) versus non-tamoxifen (p16-) treated mice. (**Fig. 4a; S5a**) We observed a significant increase in p16 tdTom^+^ primarily in immune populations following tamoxifen treatment and especially targeting alveolar macrophage population (CD45^+^ CD11b^-^ F480^+^). This cell population had a trend towards increasing frequency over time following resection (**Fig. 4b).**

**Fig. 4.**
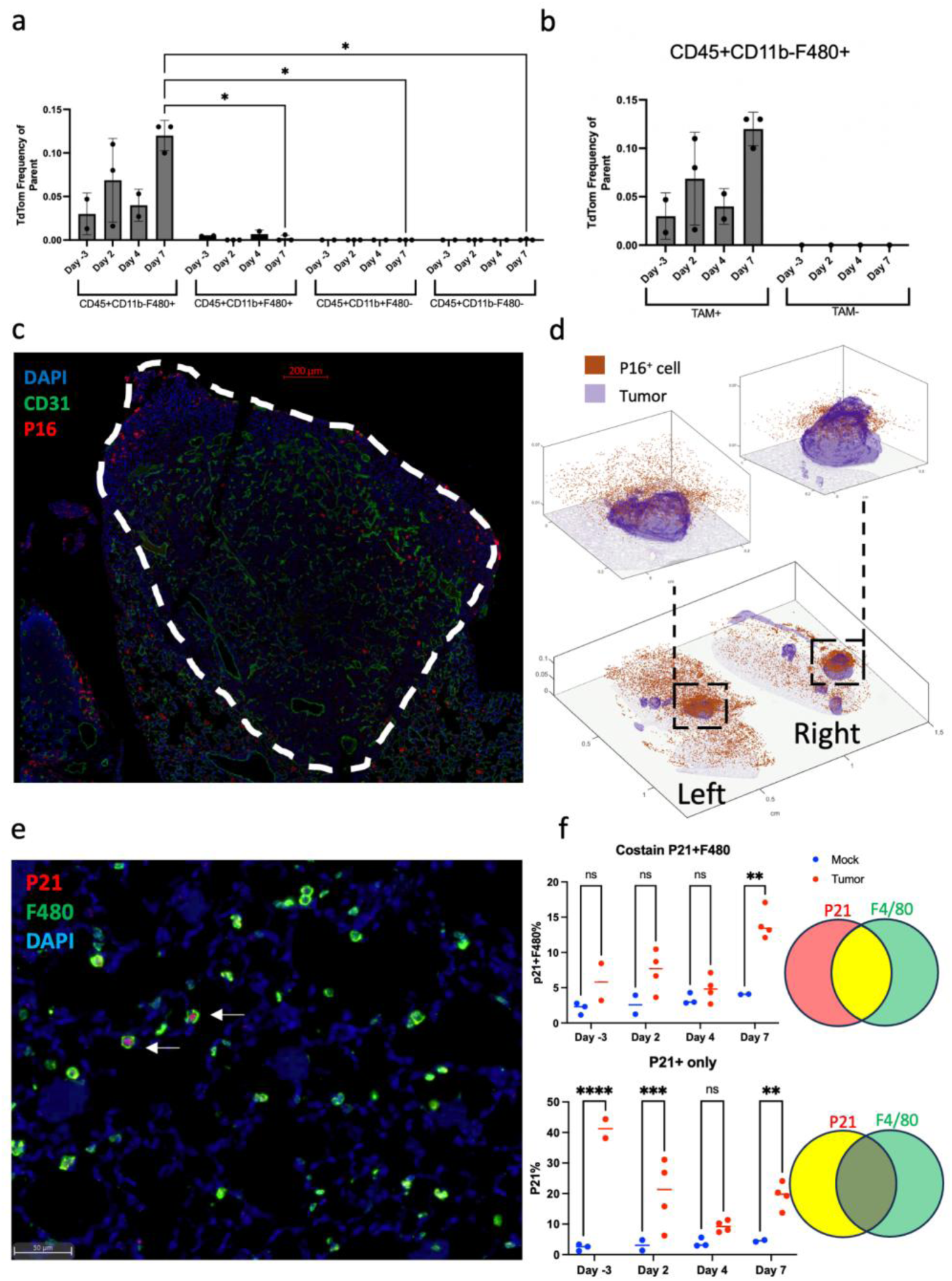
Confirmation of SnC presence in the LLC model across early timepoints. **(a)** tdTomato^+^ immune cell as frequency of parent population with comparisons between relevant immune populations. (n=2-3 per timepoint of experimental samples; n=1 tamoxifen control) **(b)** Comparing p16^+^ CD45^+^ CD11b^-^ F480^+^ to tamoxifen control (n=2-3 per timepoint of experimental samples; n=1 tamoxifen control) **(c)** Immunofluorescence staining for p16, CD31, and DAPI, showing increase of p16^+^ cell near and within metastatic LLC tumors. **(d)** 3D CODA reconstruction showing localization of p16^+^ cells from IF staining and tumors from H&E slides. **(e)** Immunofluorescence staining for p21, F480, and DAPI (arrows indicate double positive cells) **(f)** Comparison of percentage of (top) p21^+^F480^+^ cells and (bottom) p21^+^ cells across timepoints. No statistical analysis is shown for the IF images (c, e) Flow cytometry results shown as bar graphs with mean ± SD. (a, b) Mixed-effects analysis with tukey multiple comparisons test (only relevant comparisons shown) (a, b) Mixed-effects analysis with Sidak multiple comparisons test (f): * p<0.05, ** p<0.01, *** p<0.001, **** p<0.0001.

To further resolve the spatial distribution of senescent cells relative to pulmonary metastases, we performed IF staining for p16, CD31, DAPI at D9 and analyzed p16 IF staining using CODA. Compared to a naive lung, we identified a higher density of p16+ SnCs in metastatic lungs and these SnCs had a higher density in close proximity to the metastases (**Fig. 4d**)

Finaly, to confirm the accumulation of senescent alveolar macrophages in premetastatic lung (D-3,2,4,7), we performed multiplexed p21, F480, and DAPI IF staining (**Fig. 4e**) Although the p21^+^F480^+^ population was increased in the tumor bearing condition at all timepoints, it was only significant late at day 7. This result is consistent with our INKA model which demonstrated an increase in senescent alveolar macrophages with late timepoints. Interestingly, the analysis also revealed that the p21^+^F480^-^ population was significantly increased as early as three days before resection. (**Fig. 4f; S5b**).

These results suggest that the progressive accumulation of senescent alveolar macrophages associated with structural reorganization of the tissue in premetastatic lung may be delineating the local formation of PreMN.

## Discussion

There is a critical need to leverage knowledge from the earliest phases of the metastatic cascade to prevent or reduce tumor dissemination by early imaging and interception of PreMN formation.^9^ To achieve these objectives, we must understand where and when recurrences first manifest in cancer patients, essential points of cancer patient management which are not yet achieved with current clinical imaging and ctDNA approaches.^25^ Within the first two years after surgery, 80% of recurrent NSCLC patients develop recurrent disease even if neoadjuvant therapy is added.^26^ While recurrence rates differ significantly by pathological stage, distant metastases associated with early tumor cell dissemination (and probably preceding tumor resection) represent the most common pattern, with the lungs being the most favored site.^27^

Our multimodal AI-guided spatial transcriptomics in lung cancer, allowed identification in whole lung volume of a unique combination of key cell populations (alveolar macrophage), cellular processes (senescence), and structural features (ECM reorganization). Further, these stromal features, potentially associated with the morphing of PreMN, were identified much earlier than all approaches currently available in clinical settings. Braxton et al were similarly able to detect discrete PanIN lesions, otherwise challenging to be found by conventional pathology approach.^14^ We believe our study demonstrates that spatial and temporal matching of histological, molecular and imaging at a single cell resolution level, will further unlock the capture of previously undetectable stroma reorganization events (as opposed to pre-cancerous lesions or tumor-associated microenvironment) announcing subsequent metastatic risks, prior even to primary tumor resection. The concept is potentially paradigm shifting in the management of curable cancer patients but with high risk of relapse via targeting amenable stroma rather the disseminated tumor cell and intercepting the premetastatic rather the metastatic cascade.^9^

The biological pertinence of senescence and macrophages accumulation, as a hallmark of PreMN formation and metastatic risk factor, bolsters the validity of our approach. Additional biomarkers selectively associated to vasculature (lymphatic vessel structure associated Lyve1) and fibrosis (Rgcc) evoked a potential priming of the pulmonary tissue for tumor cell dissemination and survival.^22,28–30^ Past research has primarily implicated wounding in accelerating tumor growth in the acute timeframe and targeting of the wound healing cascade can reduce metastasis.^31,32^ Our present results further suggest that the chronic components of wound healing, such as senescence, angiogenesis and fibrosis may also be important in the pre-metastatic to metastatic cascade.^7,33^

In future studies we will validate these potential biomarkers of the PreMN using analysis of retrospective lung cancer resection cohorts and include them in clinical trials aimed at testing the impact of adjuvant interception of PreMN in decreasing the risk of post-resection metastatic disease.

We have indeed previously showed in clinic that the use of low dose epigenetic modifying drugs in resectable early-stage lung tumor significantly delayed the cancer recurrence and metastatic disease.^9^ In conclusion, this work serves as a proof of principle for combining a sequential timepoint analysis with 3D spatial visualization tools, such as CODA, to identify biomarkers of premetastatic stroma. This research lays the groundwork for future experiments to target specific senescent cells in metastatic models.

Our study has a number of key limitations which should be addressed. The first major limitation is the choice of timepoint for running GeoMx analysis and the limited number of samples which were run. We chose to run this analysis at day 4 timepoint due to both the tumor heterogeneity present and the availability of slides with multiple regions available for analysis. Additionally, the specific distance from tumors chosen to be analyzed as near vs far is an important caveat. It is possible that the regions around a metastasis are either completely remodeled into Pre-MN or have not yet begun to be remodeled. However, performing GSEA analysis revealed both an immune and ECM signature increased in the regions farther from the tumors. The immune signature appeared consistent with our tumor size analysis, with farther regions corresponding to more active remodeling. Finally, our findings will require further validation in retrospective cohorts to establish a significant difference in PreMN detection between patients who will develop tumor recurrence versus those without recurrence.

## Funding acknowledgement

NIH T32 MSTP grant T32GM136577; DOD LCRP HT94252410571; 2022 Johns Hopkins University Discovery Award; NIHU54CA268083.

This manuscript is the result of funding in whole or in part by the National Institutes of Health (NIH). It is subject to the NIH Public Access Policy. Through acceptance of this federal funding, NIH has been given a right to make this manuscript publicly available in PubMed Central upon the Official Date of Publication, as defined by NIH.

## Acknowledgements

We would like to acknowledge Kristen Lecksell, Elizabeth Will, and Sarah Hughes from the Oncology Tissue and Imaging Services (OTIS) Core for their assistance with histology processing and Kelly Flavahan from the Center for Infection and Inflammation Imaging (Ci3R) Core for her assistance with computed tomography imaging.

---

**Fig. S1.**
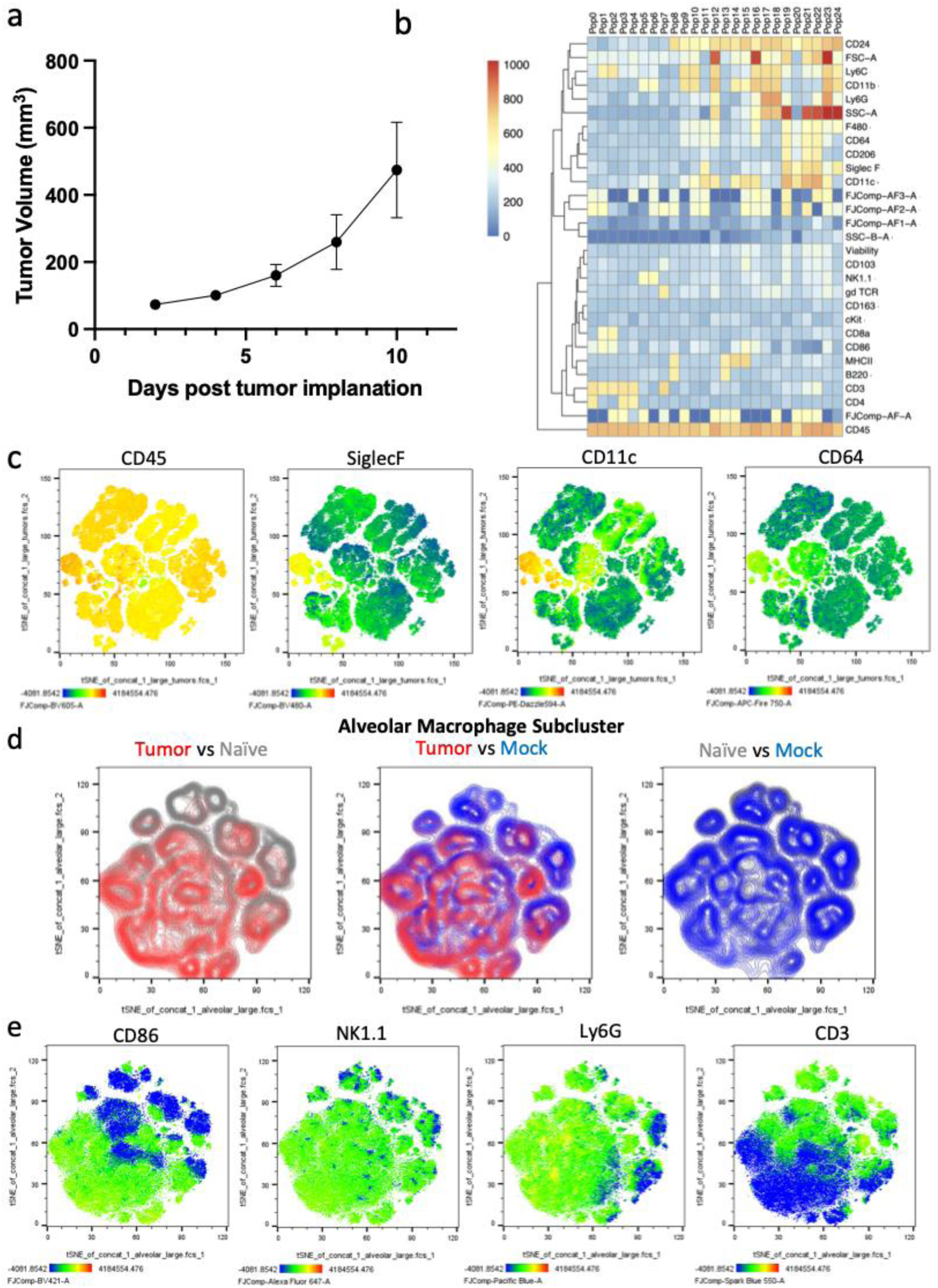
Tumor growth curve of LLC model and non-biased analysis of high parameter flow cytometry subclusters. **(a)** Tumor growth kinetics for LLC mice harvested for Pan-immune and ICS panels (n=6) **(b)** Heat map of marker expression per pan-immune flow cluster **(c)** Feature plots of CD45, siglecF, CD11c, and CD64 – alveolar macrophages indicated by arrow **(d)** Comparison of tSNE representations of the alveolar macrophage clusters between tumor, mock, and naïve groups **(e)** Feature plots of CD86, NK1.1, Ly6G, and CD3 (**Statistics**) No statistical analysis was performed for the sole tumor growth curve, heat map, or pan-immune tSNE plots (a, b, c, d, e) For tumor growth curves: Mean±SEM (a)

**Fig. S2.**
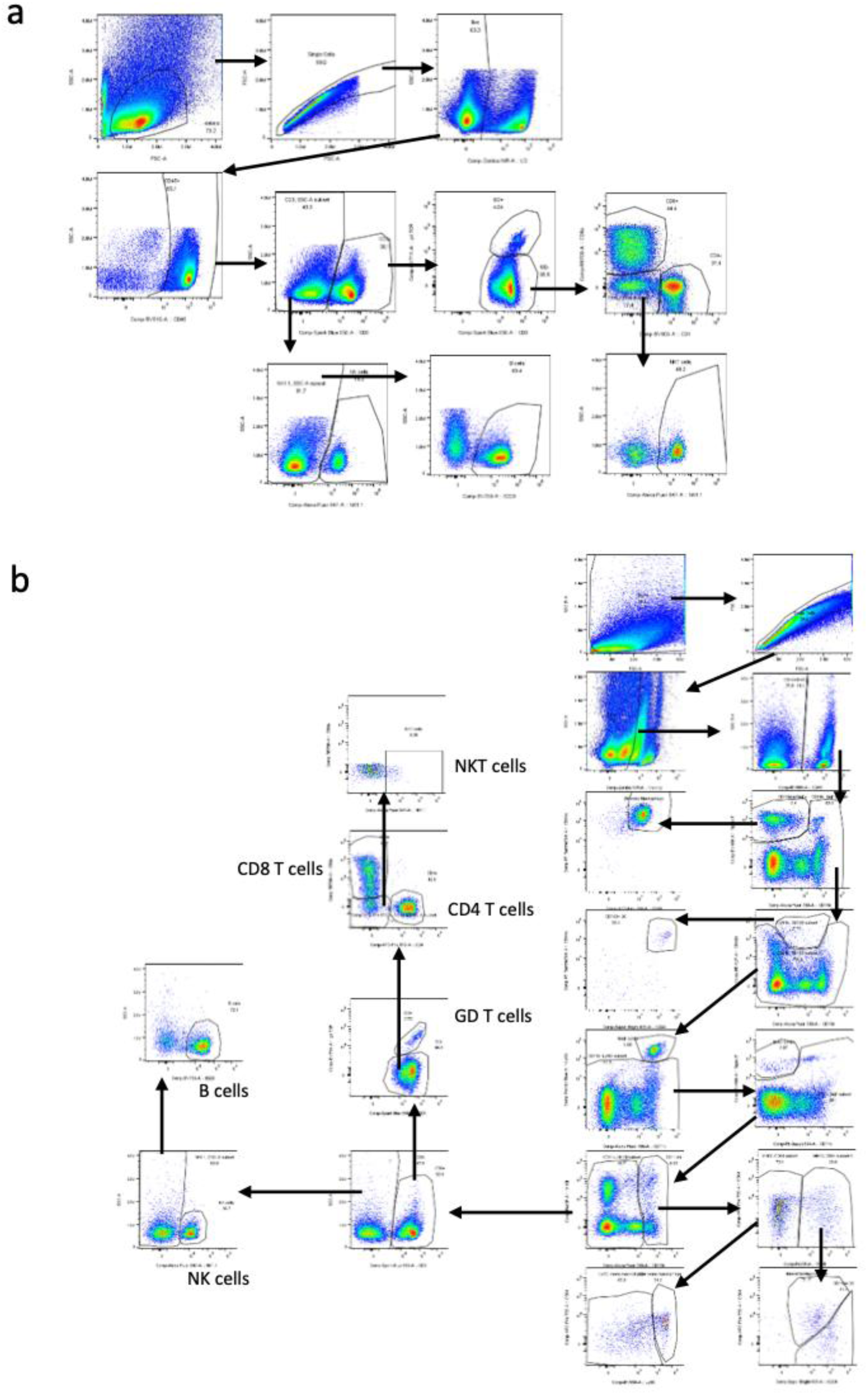
Classical flow gating schemes for high parameter flow analysis on LLC model. **(a)** ICS flow gating scheme for LLC tumor bearing, mock, and naïve mice. **(b)** Pan-immune flow gating scheme for LLC tumor bearing, mock, and naïve mice. (**Statistics**) No statistical analysis was performed for the flow gating scheme. (a, b)

**Fig. S3.**
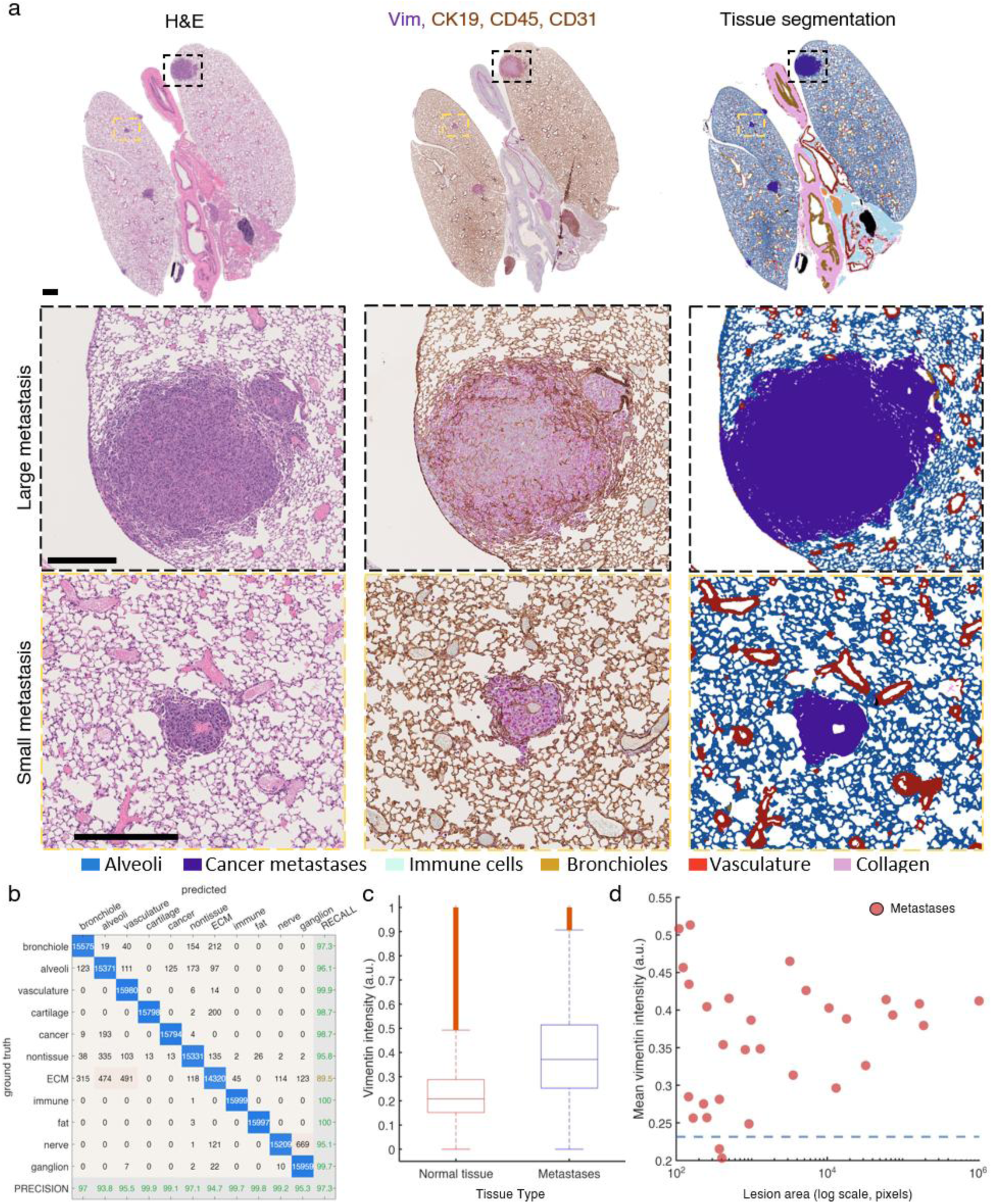
Validation of CODA segmented lung tissues and metastasis. **(a)** Validation of AI-based metastasis detection in H&E was performed using an immunohistochemistry stain performed on adjacent slides (Vimentin=red, CK19 + CD45 + CD31 = brown). Middle row shows segmentation results in large metastasis. Bottom row shows segmentation results in small metastasis. Scale bars: 5µm. **(b)** Confusion matrix showing the accuracy of tissue segmentation in H&E, with precision values indicating classification performance. **(c)** Box plot comparison of Vimentin intensity between normal tissue and metastases, showing elevated Vimentin expression in metastatic lesions. **(d)** Scatter plot of individual metastatic lesion area versus mean Vimentin intensity, showing vimentin expression across metastases of varying sizes. Dashed blue line is mean vimentin expression in normal tissue. Each point represents one metastasis.

**Fig. S4.**
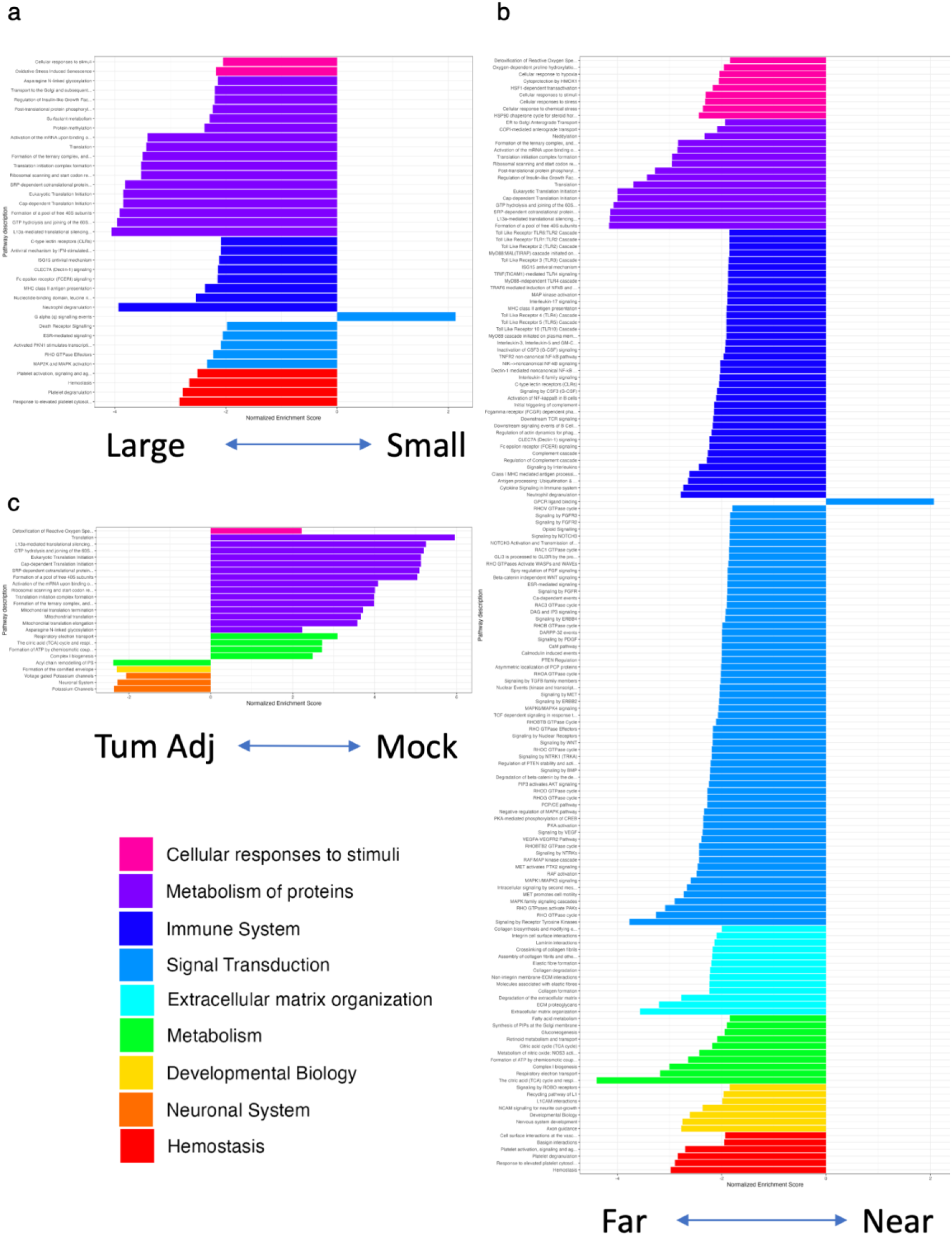
GeoMx Reactome Pathway Enrichment Plots. For all plots enriched pathways are color coded by their highest level parent pathway in Reactome **(a)** Comparison of Large versus small tumor annotations. **(b)** Comparison of Far versus Near annotations. **(c)** Comparison of tumor adjacent versus mock annotations.

**Fig. S5.**
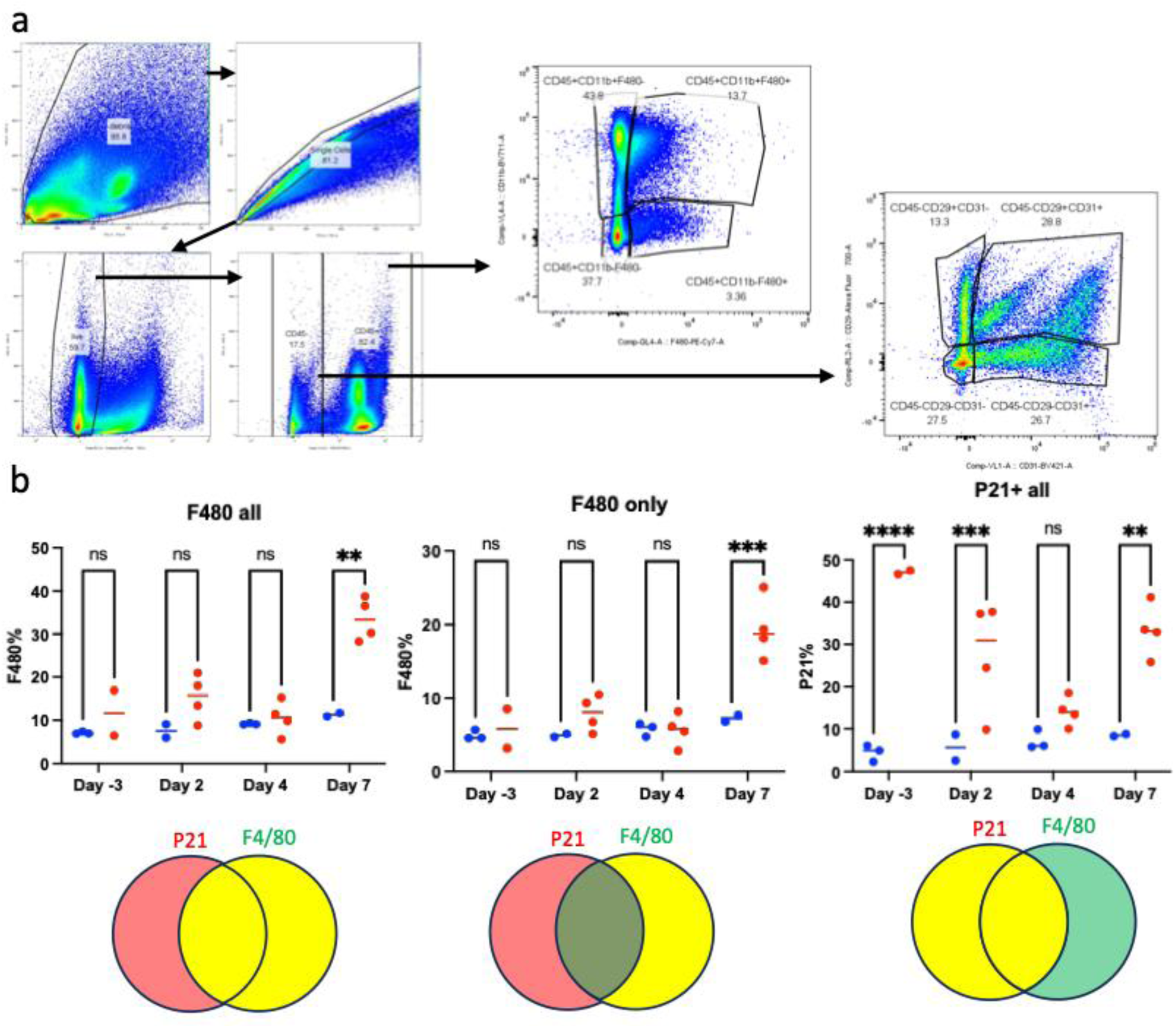
INKA Flow Gating and Additional IF Comparisons. **(a)** INKA mouse flow gating scheme. **(b)** Comparison of percentage of (left) all F480^+^ cells, (middle) only F480^+^ cells, and (right) all p21^+^ cells across timepoints. No statistical analysis is shown for the flow gating. (a) Mixed-effects analysis with Sidak multiple comparisons test (b): * p<0.05, ** p<0.01, *** p<0.001, **** p<0.0001.

## Supplemental Tables

**Supplemental Table 1.** Size distribution of lung metastases 3D mapped using CODA.

| Day | N | Total metastases | Min (µm³) | Q25 (µm³) | Median (µm³) | Mean (µm³) | Q75 (µm³) | Max (µm³) | Std Dev (µm³) | IQR (µm³) |
| --- | --- | --- | --- | --- | --- | --- | --- | --- | --- | --- |
| Day -3 | 2 | 3 | 4.15E+04 | 5.88E+04 | 1.11E+05 | 3.41E+05 | 6.81E+05 | 8.71E+05 | 4.60E+05 | 6.22E+05 |
| Day 2 | 2 | 25 | 5.36E+04 | 2.08E+05 | 4.46E+05 | 1.88E+06 | 1.37E+06 | 1.98E+07 | 4.12E+06 | 1.16E+06 |
| Day 4 | 2 | 21 | 5.53E+04 | 2.16E+05 | 5.69E+05 | 1.09E+07 | 4.66E+06 | 1.32E+08 | 3.01E+07 | 4.44E+06 |
| Day 7 | 2 | 78 | 3.63E+04 | 3.11E+05 | 2.73E+06 | 4.56E+07 | 2.81E+07 | 6.04E+08 | 1.08E+08 | 2.78E+07 |

**Supplemental Table 2.** Pan-immune flow cytometry panel, which was run on an aurora flow cytometer.

| Fluorophore | Antigen | Clone | Dilution | Manufacturer |
| --- | --- | --- | --- | --- |
| BV421 | CD86 | GL-1 | 200 | Biolegend |
| SuperBright436 | CD24 | M1/69 | 300 | ThermoFisher |
| Pacific Blue | Ly6g | 1A8 | 250 | Biolegend |
| BV480 | Siglec-F | E50-2440 | 100 | BD Biosciences |
| BV605 | CD45 | 30-F11 | 300 | Biolegend |
| BV650 | Ly6C | HK1.4 | 1200 | Biolegend |
| BV711 | γΔ TCR | GL3 | 200 | BD Biosciences |
| BV750 | B220 | RA3-6B2 | 200 | Biolegend |
| BV785 | F4/80 | BM8 | 300 | Biolegend |
| AF488 | cKit | 2B8 | 200 | Biolegend |
| SparkBlue550 | CD3 | 17A2 | 100 | Biolegend |
| PerCP | MHC-II | M5/114.15.2 | 200 | Biolegend |
| BB700 | CD8a | 53-6.7 | 100 | BD Biosciences |
| PE | CD163 | S15049I | 300 | Biolegend |
| PE-Dazzle594 | CD11c | N418 | 700 | Biolegend |
| PE-Cy7 | CD103 | QA17A24 | 100 | Biolegend |
| APC | CD206 | C068C2 | 200 | Biolegend |
| AF647 | NK1.1 | PK136 | 200 | Biolegend |
| AF700 | CD11b | M1/70 | 200 | Biolegend |
| Zombie NIR | Viability | - | 3000 | Biolegend |
| APC-Fire750 | CD64 | X54-5/7.1 | 100 | Biolegend |
| APC-Fire810 | CD4 | GK1.5 | 100 | Biolegend |

**Supplemental Table 3.**
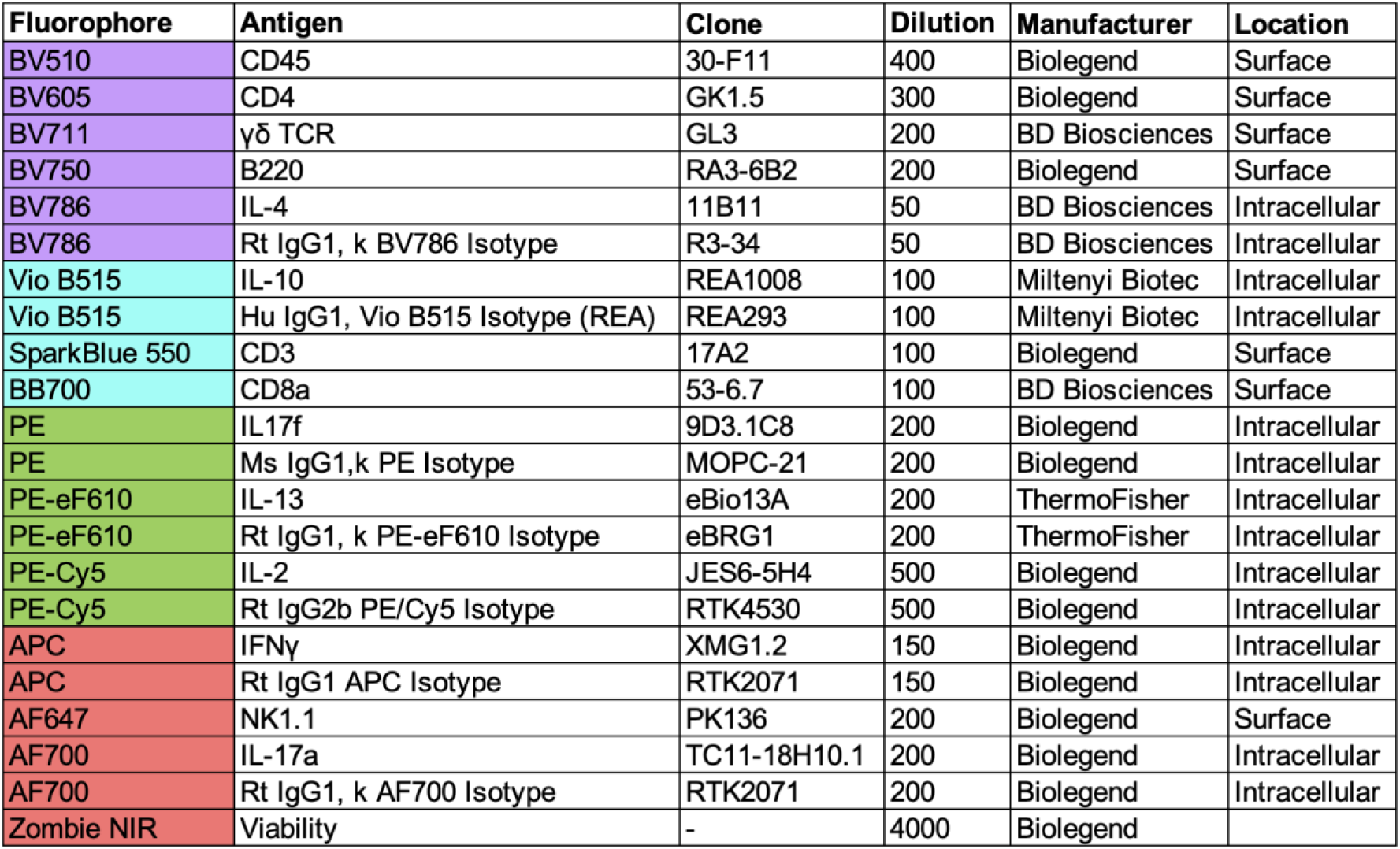
Intracellular Cytokine flow cytometer panel, which was run on an aurora flow cytometer.

| Fluorophore | Antigen | Clone | Dilution | Manufacturer | Location |
| --- | --- | --- | --- | --- | --- |
| BV510 | CD45 | 30-F11 | 400 | Biolegend | Surface |
| BV605 | CD4 | GK1.5 | 300 | Biolegend | Surface |
| BV711 | γδ TCR | GL3 | 200 | BD Biosciences | Surface |
| BV750 | B220 | RA3-6B2 | 200 | Biolegend | Surface |
| BV786 | IL-4 | 11B11 | 50 | BD Biosciences | Intracellular |
| BV786 | Rt IgG1, k BV786 Isotype | R3-34 | 50 | BD Biosciences | Intracellular |
| Vio B515 | IL-10 | REA1008 | 100 | Miltenyi Biotec | Intracellular |
| Vio B515 | Hu IgG1, Vio B515 Isotype (REA) | REA293 | 100 | Miltenyi Biotec | Intracellular |
| SparkBlue 550 | CD3 | 17A2 | 100 | Biolegend | Surface |
| BB700 | CD8a | 53-6.7 | 100 | BD Biosciences | Surface |
| PE | IL17f | 9D3.1C8 | 200 | Biolegend | Intracellular |
| PE | Ms IgG1,k PE Isotype | MOPC-21 | 200 | Biolegend | Intracellular |
| PE-eF610 | IL-13 | eBio13A | 200 | ThermoFisher | Intracellular |
| PE-eF610 | Rt IgG1, k PE-eF610 Isotype | eBRG1 | 200 | ThermoFisher | Intracellular |
| PE-Cy5 | IL-2 | JES6-5H4 | 500 | Biolegend | Intracellular |
| PE-Cy5 | Rt IgG2b PE/Cy5 Isotype | RTK4530 | 500 | Biolegend | Intracellular |
| APC | IFNγ | XMG1.2 | 150 | Biolegend | Intracellular |
| APC | Rt IgG1 APC Isotype | RTK2071 | 150 | Biolegend | Intracellular |
| AF647 | NK1.1 | PK136 | 200 | Biolegend | Surface |
| AF700 | IL-17a | TC11-18H10.1 | 200 | Biolegend | Intracellular |
| AF700 | Rt IgG1, k AF700 Isotype | RTK2071 | 200 | Biolegend | Intracellular |
| Zombie NIR | Viability | - | 4000 | Biolegend |  |

**Supplemental Table 4.** Lewis lung carcinoma (LLC) INKA flow panel, which was run on an attune flow cytometer.

| Fluorophore | Antigen | Clone | Dilution | Manufacturer |
| --- | --- | --- | --- | --- |
| BV421 | CD31 | 390 | 200 | BioLegend |
| BV605 | CD45 | 30-F11 | 150 | BioLegend |
| BV711 | CD11b | M1/70 | 300 | BioLegend |
| AF488 | EpCAM | 17A2 | 50 | BioLegend |
| tdTomato | p16 | - | - | - |
| PE-Cy7 | F4/80 | BM8 | 250 | BioLegend |
| AF647 | CD146 | ME-9F1 | 2000 | BioLegend |
| AF700 | CD29 | HMβ1-1 | 100 | BioLegend |
| eFluor 780 | Viability | - | 1000 | ThermoFisher |

## Materials and Methods

### Mice

All procedures were performed in compliance with approved Johns Hopkins Animal Care and Use Committee (ACUC) protocols. Mice were cared for in the Bunting Blaustein and Koch Cancer Research Buildings. Unless otherwise mentioned, female C57BL/6J mice were used for tumor stock expansion and all experiments. The only exception was the use of the p16-CreERT2;Ai14 mice (INKA mouse model) as a reporter for p16 reporter, which were previously developed and characterized.^24^

### Tamoxifen Induction of INKA mouse model

To induce the INKA model (described above) to express tdTomato signal in cells with p16 promoter activity 5 days of tamoxifen (70 mg/kg/day) were given 7 days in advance of tissue harvest. Tamoxifen (Sigma-Aldrich) was solubilized in corn oil (Millipore Sigma) (14mg tamoxifen/ 1ml corn oil) by allowing the mixture to sit overnight at 37C in a rotating mixer. 5ul was injected IP per gram of mouse using a 26G TB syringes.^24^

### Lewis Lung Carcinoma (LLC) implantation surgery

Prior to all surgeries mice were anesthetized, shaved, and the surgical site was sterilized. The LLC tumors experienced at least 2 passages in vivo prior to implantation in experimental mice to ensure proper growth kinetics. For all procedures animals were allowed to recover on a heated surface, returned to their cages, and observed for pain after 24 hours for delivery of Carprofen (4-5 mg/kg) (MWI Animal Health) subcutaneously.

For the initial implantation a 1 cm incision was made in the flank of the mouse and then a 3-4 mm^3^ tumor tissue was subcutaneously transplanted by tweezer. This tumor was either sourced from our liquid nitrogen stock or from another mouse. Finally, either sterile silk 4-0 sutures or surgical staples will be used to close the wounds. Tumors were monitored three times a week using a Digital Caliper (Kynup) to measure the length and width of each tumor. Tumor volume was calculated as ½ x shortest side x longest side x longest side. After 9-12 days, most of the tumors reach 300-700 mm^3^ and resection is performed when the flank tumor reaches a burden of 375-625 mm^3^. Mice were distributed into groups to minimize differences in average volume and standard deviation of volume between group. These groups were then randomized to treatment conditions.

For the resection surgery, the skin was incised 1cm outside the border of the tumor. Total tumor resection was carried out and sterile silk 4-0 sutures or surgical staples were used to close the wounds. Mice were monitored closely following resection for euthanasia criteria, primary tumor regrowth, weight loss, and pain. Resected tumors were either discard, implanted into a mouse for a subsequent passage, or frozen to generate tumor stock.

### Computed Tomography (CT) Scanning

Live imaging with cancer recurrence and progression was performed in Koch Cancer Research Building by personnel trained in handling small animals/rodents. The mice were anesthetized with isoflurane for the entire duration of the imaging and received both heat support and ophthalmic ointment. All personnel handling the rodents wore sterile gloves and masks throughout the procedure. NanoScan PC (Mediso) was used to scan the mice with the following parameters, Image Type: DERIVED\PRIMARY\AXIAL; Software Versions: VivoQuant 2020patch1-build3; Study Description: Factory/Clinical Protocols/CT/CT Reconstruction/High Resolution Reconstruction; Series Description: CTRecon,Post Reconstruction,FBP,V40,Sl40,F:Ram,C100,M30,S0; Slice Thickness: 0.04 mm; Tube voltage: 50 kVp; Spacing Between Slices: 0.04 mm; Software Versions: Nucline-3.04.012.0000; Exposure Time: 300 s; X-Ray Tube Current: 1, Exposure in µAs: 201; Filter Type: NONE; Position: HFP; Revolution Time: 432.0; Single Collimation Width: 31.2 µm; Total Collimation Width: 30346.4 µm; Photometric Interpretation: MONOCHROME2; Pixel Spacing: 0.04065\0.04065; Units: Hounsfield unit. Images were read by Dr. Cheng Ting Lin, MD for presence of metastasis.

### Tissue Processing for Flow Cytometry

#### Lung Processing for Both Aurora Panels

Mouse lungs were harvested and dissected into single lobes with care taken to separate and remove the trachea, heart, thymus, and adjacent lymph nodes. The lungs were rinsed once with DPBS (Gibco - without calcium, magnesium, or phenol red) and placed into a seperate gentleMACS™ tubes (Miltenyi Biotec) containing 4.02ml of pre-chilled RPMI 1640 medium without HEPES and with L-glutamine and Phenol Red (Gibco) supplemented additionally with 0.1% L-glutamine 200mM (Quality Biological), 0.1% HEPES buffer 1M (gibco), and 5% heat-inactivated FBS (GeminiBio). After harvesting, 160ul of 20,000 Mandl U/mL Liberase TM (Sigma) and 820ul of 1mg/ml DNAse1 (Sigma) were quickly added to each tube. The tubes were then connected to a gentleMACS™ dissociator (Miltenyi Biotec) and the 37C_m_LDK_1 program was run to dissociate and digest the tissue. Following digestion, 492ul of 100 mM EDTA 7.2 pH was added to each tube to neutralized the digestion. The 100mM EDTA was prepared by combining 100ml 0.5M EDTA (invitrogen) and 500ml PBS (gibco) and adjusting the pH to 7.2 with HCl (Avantor Sciences). This digest was added into a 100um cell strainer (Falcon) in a 50ml conical tube on ice and the flat end of a syringe plunger was used to grind the tissue through the filter. This strainer was washed twice with the aforementioned RPMI media supplemented with an additional 0.4% of 0.5M EDTA. The syringe plunger was used to mash the filter after every wash. The liquid that had passed through the filter was centrifuged at 500g for 5 mins at 4C to generate a pellet, which was resuspended in DPBS after carefully aspirating. This cell suspension was then split in half, as this was the last processing step in common between the pan-immune and cytokine panels (described in below sections).

#### Lung Processing Specific for Pan-Immune Aurora Panel

The portion of the cell suspension that was proportioned to pan-immune panel was centrifuged at 500g for 5 mins at 4C to generate a pellet. This pellet was resuspended with 2.5ml of ACK lysis buffer (Quality Biological) and incubated at room temperature for 5 mins. 30ml of DPBS was subsequently added to prevent further lysis and the tube was again centrifuged to generate a pellet. This pellet was resuspended, filtered once through a 100um cell strainer (Falcon) to remove any RBC debris. Cell counting was then performed using ViaStain AOPI Staining Solution (Revvity) imaged on a Cellometer K2 (Nexcelom). The staining procedure for the pan-immune panel is described in depth in the next section.

#### Lung Processing Specific for Cytokine Aurora Panel

The portion of the cell suspension that was proportioned to cytokine panel was run through a Percoll gradient to remove RBCs and stromal cells. The day before the harvest the four Percoll dilutions were generated using Percoll (GE Healthcare), 10x PBS (gibco), 1x PBS (gibco), RPMI (gibco): 100% Percoll (1:9, 10x PBS to Percoll), 80% Percoll (2:8, 1xPBS to 100% Percoll), 40% Percoll (1:1, 70ml RPMI to 80% Percoll), and 20% (2:8, 100% Percoll to 1xPBS). All Percoll dilutions were brought to room temperature before use in gradient. Each cell suspension was centrifuged at 500g for 5 mins at 4C to generate a pellet, which was then resuspend with 4ml 80% Percoll and transferred to a 15ml conical tube. 4ml 40% Percoll was then slowly laid on top of the 80% layer using a serological pipette with care taken to not disturb the previous layer. This procedure was repeated with 3ml of 20% Percoll. The subsequent gradients were centrifuged at 2,100g for 30mins at room temperature with accelerators and breaks set to the lowest value. Once complete the top debris layer (approximately to the 8ml mark) was aspirated, and the 80-40 interface containing the immune cells (approximately between the 7ml and 1.5ml mark) were pipetted into a new 15ml conical tube. This transferred Percoll was then diluted with 8ml of 1x DPBS, shaken well, and centrifuged at 500g for 5 mins at 4C. The pellet was resuspended and cell counting was then performed using ViaStain AOPI Staining Solution (Revvity) imaged on a Cellometer K2 (Nexcelom). The staining procedure for the cytokine panel is described in depth in the next section.

#### Lung Processing for Attune Panel

Mouse lungs were harvested and dissected into single lobes with care taken to separate and remove the trachea, heart, thymus, and adjacent lymph nodes. Each mouse’s lungs were placed in separate 100mm petri dishes containing 5ml RPMI 1640 medium without HEPES and with L-glutamine and Phenol Red (Gibco) supplemented additionally with 0.1% L-glutamine 200mM (Quality Biological), 0.1% HEPES buffer 1M (gibco), and 5% heat-inactivated FBS (GeminiBio). Lungs were diced using razor blades (< 1mm pieces) were transferred into 15ml conical tubes. Digestion was performed by adding 100ul of 20,000 Mandl U/mL Liberase TM (Sigma) and 1ml of 1mg/ml DNAse1 (Sigma) and allowing the tubes to incubate for 25mins at 37C. To neutralize the digestion 600ul of 100 mM EDTA 7.2 pH was added to each tube. The 100mM EDTA was prepared by combining 100ml 0.5M EDTA (invitrogen) and 500ml PBS (gibco) and adjusting the pH to 7.2 with HCl (Avantor Sciences). This digest was added into a 100um cell strainer (Falcon) in a 50ml conical tube on ice and the flat end of a syringe plunger was used to grind the tissue through the filter. This strainer was washed twice with the aforementioned RPMI media supplemented with an additional 0.4% of 0.5M EDTA. The syringe plunger was used to mash the filter after every wash. The liquid that had passed through the filter was centrifuged at 500g for 5 mins at 4C to generate a pellet. This pellet was resuspended with 5ml of ACK lysis buffer (Quality Biological) and incubated at room temperature for 5 mins. 30ml of RPMI+EDTA media was subsequently added to prevent further lysis and the tube was again centrifuged to generate a pellet. This pellet was resuspended, filtered once through a 100um cell strainer (Falcon) to remove any RBC debris. Cell counting was then performed using ViaStain AOPI Staining Solution (Revvity) imaged on a Cellometer K2 (Nexcelom). The staining procedure for the attune panel is described in depth in the next section.

### Multi-Parameter Flow Cytometry

#### Pan-Immune Aurora Panel

After the final wash, resuspend cells in DPBS were plated at 500k cells/ml in the appropriate wells of a 96 well plate. Plates were centrifuge 400 - 500 x G for 5 minutes, decanted, and cell pellets were washed with 200ul of DPBS. Cells were then stained with zombie Near IR viability stain dye (Biolegend) at a 1:3000 dilution in DPBS for 30 minutes, covered, on ice. Unstained, Single Stain Control (SSC), and Zombie NIR Fluorescence Minus One (FMO) conditions were instead incubated with plain DPBS. Following incubation in Viability Buffer cells were washed twice using flow wash buffer: DPBS + 1% Bovine Serum Albumin (BSA) (Sigma) + 1mM EDTA (Invitrogen). For all aurora panels FACS buffer was supplemented with 1mM L-Ascorbic acid (Sigma Aldrich). Samples were then stained with the appropriate antibody cocktail for 45 minutes, covered, on ice. All antibody cocktails contained 1:50 TruStain FcX™ (anti-mouse CD16/32) Antibody (Biolegend) + 1:20 True-Stain Monocyte Blocker™ (Biolegend) + 1:20 Super Bright Complete Staining Buffer (eBioscience). All stain samples, including all experimental conditions, were stained with the antibodies in the following table at the indicated dilution (**Table S2**). SSC controls and FMOs included either only one antibody or all but one antibody, respectively. The volume of each cocktail was increased to 100ul after accounting for the volume of antibodies, Fc blocker, monocyte blocker, and super bright buffer. Following incubation, cells were washed twice with wash buffer and fixed with FluoroFix™ Buffer (Biolegend) for 15 minutes, covered, at room temperature. Cells were washed twice with wash buffer and run on a spectral flow cytometer (Cytek Aurora) the following day.

#### Preparation of Ultracomp beads

0.5ul of appropriate antibody were added to 1-2 drops of ultracomp beads and incubated for 7 min in the dark at room temperature. Unstained ultracomp beads received no antibodies. 500ul of DPBS were added to each condition.

#### Cytokine Aurora Panel

After the final wash, resuspend cells in DPBS were plated at 500k cells/ml in the appropriate wells of a 96 well plate. To create a 1x stimulation media, Iscove’s Modified Dulbecco’s Medium (IMDM) (with L-glutamine and without Phenol Red, α-thioglycerol, 2-mercaptoethanol) was supplemented with 0.2% Cell Stimulation Cocktail Plus Protein Transport Inhibitors (Bioscience) and 5% heat-inactivated FBS (ThermoFischer). Each well was resuspend 2.5x10^6 cells/mL with the stimulation media and left in an incubator for 4-6 hours at 37C. Three times plates were centrifuged at 500xg for 5 minutes at 4°C, decanted, and resuspended in 200ul of chilled DPBS. Cells were then stained with zombie Near IR viability stain dye (Biolegend) at a 1:4000 dilution in DPBS for 30 minutes, covered, on ice. Unstained, SSC, and Zombie NIR Fluorescence Minus One (FMO) conditions were instead incubated with plain DPBS. Samples were then stained with the appropriate surface antibody cocktails for 45 minutes, covered, on ice. All antibody cocktails contained 1:50 TruStain FcX™ (anti-mouse CD16/32) Antibody (Biolegend) + 1:20 True-Stain Monocyte Blocker™ (Biolegend) + 1:20 Super Bright Complete Staining Buffer (eBioscience). All stain samples, including all experimental conditions, were stained with the surface antibodies in the following table at the indicated dilution (**Table S3**). SSC controls and FMOs included either only one antibody or all but one antibody, respectively. Cells were washed once with FACS buffer and incubated with Cyto-Fast™ Fix (Biolegend) for 20mins at room temperature. For all aurora panels FACS buffer was supplemented with 1mM L-Ascorbic acid (Sigma Aldrich). Wells were washed twice with P/W buffer which was composed of 10% Perm Wash (Biolegend) with 90% DI H2O. Intracellular antibody cocktails were generated in a similar manner as the surface antibody cocktails with the a few notable exceptions: P/W buffer was used instead of FACS buffer and FMOs contained isotype antibodies instead of no antibody. Cells were stained with the intracellular cocktails for 30 mins at room temperature. Samples were then washed once with P/W and FACS buffer and resuspend with FACS buffer for overnight storage at 4C. Cells were washed twice with wash buffer and run on a spectral flow cytometer (Cytek Aurora) the following day.

#### Attune INKA Flow Panels

After the final wash, cells were resuspended in DPBS and plated at 500k cells/ml for lungs and 1million cells/ml for tumors in the appropriate wells of a 96 well plate. INKA mice given tamoxifen were used for experimental, tdTomato SSC, and all FMOs except for the tdTomato FMO. INKA mice not given tamoxifen were used for all other conditions. Plates were centrifuge 400 - 500 x G for 5 minutes, decanted, and cell pellets were washed with 200ul of DPBS. Cells were then stained with eflouro 780 Viability Stain (eBioscience) at a 1:1000 dilution in DPBS for 30 minutes, covered, on ice. Unstained, Single Stain Control (SSC), and Zombie NIR Fluorescence Minus One (FMO) conditions were instead incubated with plain DPBS. Following incubation in Viability Buffer cells were washed twice using FACS buffer: DPBS + 1% Bovine Serum Albumin (BSA) (Sigma) + 1mM EDTA (Invitrogen). Samples were then stained with the appropriate antibody cocktail for 45 minutes, covered, on ice. All antibody cocktails contained 1:50 TruStain FcX™ (anti-mouse CD16/32) Antibody (Biolegend) + 1:20 True-Stain Monocyte Blocker™ (Biolegend) + 1:20 Super Bright Complete Staining Buffer (eBioscience). All stain samples, including all experimental conditions, were stained with the antibodies in the following table at the indicated dilution (**Table S4**). SSC controls and FMOs included either only one antibody or all but one antibody, respectively. The volume of each cocktail was increased to 100ul after accounting for the volume of antibodies, Fc blocker, monocyte blocker, and super bright buffer. Following incubation, cells were washed twice with wash buffer and fixed with FluoroFix™ Buffer (Biolegend) for 15 minutes, covered, at room temperature. Cells were washed twice with wash buffer and run on a conventional flow cytometer (Attune NXT in the SKCCC Flow Cytometry Technology Development Center) the following day.

### t-distributed stochastic neighbor embedding (tSNE) flow analysis

All full stained, experimental conditions were analyzed by tSNE on FlowJo. Samples were gated to reveal population of interest, either all immune cells (live+CD45+) or alveolar macrophages (CD45^+^ CD11b^mid-low^ SigF^+^ CD11c^+^ CD64^+^). Subsequently, QC was performed using the flowAI program.^34^ Briefly, all compensation + time channels were included, Anomalies to exclude = All checks, Second fraction FR = 0.1, Alpha FR = 0.01, maximum changepoints = 3, changepoint penalty = 200, Dynamic range check side = both, Decompose flow rate = TRUE, Clean up working dir = TRUE. Down-sampling was then performed, such that all samples contributed the same number of cells to the analysis. The immune cell samples were down-sampled to 10,000 events and the alveolar macrophage samples were down-sampled to 4,500 events. These samples were then concatenated to make a single flow file: Format: FCS3, Include events = Include all, Parameters = all compensated parameters, Group concatenation = Concatenate all files together, Additional parameters = experimental group + animal number, Spread distribution of keyword data = true. tSNE was then with the following distinction, the immune group used all compensation (except for CD45+ and Live/Dead which had already been gated), FSC-A, and SSC-B-A; and the alveolar macrophage group used all compensation (except for CD45+, SigF+, CD11c, CD64, and Live/Dead which had already been gated), FSC-A, and SSC-B-A. The following parameters were used for tSNE: Learning configuration = Auto (opt-SNE), iterations = 1000, perplexity = 30, KNN algorithm = Exact (vantage point tree), gradient algorithm = Barnes-Hut. Flow SOM (3.0.18 PP) was then run with the same parameter limitations as tSNE analysis with number of meta clusters equal to 25 for the immune population and 8 for the alveolar macrophage population.^35^ The following parameters were then used for flowSOM: Apply on map = None, SOM grid size (W x H) = 10 x 10, view layout = Minimum spanning tree, Plot channel = all as piecharts, Node scale = 100%, Background metaclusters = TRUE, create PDF = TRUE, Save the R script and output messages = TRUE. At this point, the overall tSNE and feature plots were generated using FlowJo. For the group specific comparisons, all groups per condition were down sampled, such that they had the same number of cells. The immune populations were downsampled to 60,000 events and the alveolar macrophage populations were downsampled to 27000 events. Contour plots were then generated by comparing two groups at a time with the contour level parameter = 2%.

### Serial Histological Sectioning

The procedure for harvesting mouse lungs has been previously described. In brief lungs were harvested by first opening the mouse’s sternum, inserting a catheter into an incision in the trachea, and slowly injecting approximately 1ml of Neutral buffer formalin (Sigma) with a tuberculin syringe 25G5/8 (BD), so as to not over or under inflate the lungs. A knot was tied below the incision to keep the NBF in the lungs and the entire lung was removed en bloc.^36^ The lungs were left in a 50ml conical tube containing 40ml of NBF was left for 2 days on an orbital rocker. Subsequently, the lungs were washed with PBS and moved into a 70% ethanol solution. The lungs were then given to the Johns Hopkins Oncology Tissue and Imaging Services (OTS) core to perform dehydration and embedding.

For the mock treated lungs the following sectioning approach was utilized: 200 four µm thick sections were taken with every third section stained for H&E, every other third section left unstained, and every other third section discarded. For the tumor bearing lungs the following sectioning approach was utilized: 300 sections were taken with every third section stained for H&E, every other third section left unstained, and every other third section also left unstained. All blocks were sectioned at a thickness of 4um with approximately 1.2mm of the block consumed in the process.

We sectioned two blocks at each timepoint. For the time points which contained mice with and without tumors detected on CT (day 4 and 7), 1 of each was chosen.

### Histological staining

#### Hematoxylin and eosin (H&E) staining

H&E staining was performed by the Johns Hopkins Oncology Tissue and Imaging Services (OTIS) core of Johns Hopkins University on formalin-fixed, paraffin-embedded (FFPE) tissue sections using a Leica ST5020 auto-stainer.

#### CD45+ stain

Immunostaining was performed at the OTIS core. Immunolabeling for CD45 was performed on formalin-fixed, paraffin embedded sections on a Ventana Discovery Ultra autostainer (Roche Diagnostics). Briefly, following dewaxing and rehydration on board, epitope retrieval was performed using Ventana Ultra CC1 buffer (catalog# 6414575001, Roche Diagnostics) at 95°C for 64 minutes. Primary antibody, anti-CD45 (1:200 dilution; catalog# 70257S, Cell Signaling Technologies) was applied at 36°C for 60 minutes. Primary antibodies were detected using an anti-rabbit HQ detection system (catalog# 7017936001 and 7017812001, Roche Diagnostics) followed by Chromomap DAB IHC detection kit (catalog # 5266645001, Roche Diagnostics), counterstaining with Mayer’s hematoxylin, dehydration and mounting.

#### CK19+ stain

Immunostaining was performed at the OTIS core. Immunolabeling for CK19 was performed on formalin-fixed, paraffin embedded sections on a Ventana Discovery Ultra autostainer (Roche Diagnostics). Briefly, following dewaxing and rehydration on board, epitope retrieval was performed using Ventana Ultra CC1 buffer (catalog# 6414575001, Roche Diagnostics) at 100°C for 64 minutes. Primary antibody, anti-CK19 (1:800 dilution; catalog# ab133496, Abcam) was applied at 36°C for 60 minutes. Primary antibodies were detected using an anti-rabbit HQ detection system (catalog# 7017936001 and 7017812001, Roche Diagnostics) followed by Chromomap DAB IHC detection kit (catalog # 5266645001, Roche Diagnostics), counterstaining with Mayer’s hematoxylin, dehydration and mounting.

#### Vimentin+ stain

Immunostaining was performed at the at the OTIS core. Immunolabeling for Vimentin was performed on formalin-fixed, paraffin embedded sections on a Ventana Discovery Ultra autostainer (Roche Diagnostics). Briefly, following dewaxing and rehydration on board, epitope retrieval was performed using Ventana Ultra CC1 buffer (catalog# 6414575001, Roche Diagnostics) at 95°C for 64 minutes. Primary antibody, anti-Vimentin (1:100 dilution; catalog# 5741S, Cell Signaling Technologies) was applied at 36°C for 60 minutes. Primary antibodies were detected using an anti-rabbit HQ detection system (catalog# 7017936001 and 7017812001, Roche Diagnostics) followed by Chromomap DAB IHC detection kit (catalog # 5266645001, Roche Diagnostics), counterstaining with Mayer’s hematoxylin, dehydration and mounting.

#### CD31+CD45+CK19+Vimentin stain

Immunostaining was performed at the at the OTIS core. Immunolabeling for CD31+CD45+CK19+Vimentin detection was performed on formalin-fixed, paraffin embedded sections on a Ventana Discovery Ultra autostainer (Roche Diagnostics). Briefly, following dewaxing and rehydration on board, epitope retrieval was performed using Ventana Ultra CC1 buffer (catalog# 6414575001, Roche Diagnostics) at 96°C for 64 minutes. Primary antibody, anti-CD31 (1:2000 dilution; catalog# ab182981, Abcam) was applied at 36°C for 3 hours and 20 minutes. CD31 primary antibodies were detected using an anti-rabbit HQ detection system (catalog# 7017936001 and 7017812001, Roche Diagnostics) followed by Chromomap DAB IHC detection kit (catalog # 5266645001, Roche Diagnostics). Following CD31 detection, primary and secondary antibodies from the first round of staining were stripped on board using Ventana Ultra CC1 buffer at 95°C for 12 minutes. Primary antibody, anti-CD45 (1:200 dilution; catalog# 70257S, Cell Signaling Technologies) was applied at 36°C for 60 minutes. CD45 primary antibodies were detected using an anti-rabbit HQ detection system (catalog# 7017936001 and 7017812001, Roche Diagnostics) followed by Chromomap DAB IHC detection kit (catalog # 5266645001, Roche Diagnostics). Following CD45 detection, primary and secondary antibodies from the second round of staining were stripped on board using Ventana Ultra CC1 buffer at 95°C for 12 minutes. Primary antibody, anti-CK19 (1:800 dilution; catalog# ab133496, Abcam) was applied at 36°C for 60 minutes. CK19 primary antibodies were detected using an anti-rabbit HQ detection system (catalog# 7017936001 and 7017812001, Roche Diagnostics) followed by Chromomap DAB IHC detection kit (catalog # 5266645001, Roche Diagnostics). Following CK19 detection, primary and secondary antibodies from the third round of staining were stripped on board using Ventana Ultra CC1 buffer at 95°C for 12 minutes. Primary antibody, anti-Vimentin (1:100 dilution; catalog# 5741S, Cell Signaling Technologies) was applied at 36°C for 60 minutes. Vimentin primary antibodies were detected using an anti-rabbit HQ detection system (catalog# 7017936001 and 7017812001, Roche Diagnostics) followed by Discovery Purple Detection kit (catalog# 7053983001, Roche Diagnostics), counterstaining with Mayer’s hematoxylin, dehydration and mounting.

#### Slide scanning

All histologically stained sections were imaged using a Hamamatsu NanoZoomer S210 bright field slide scanning microscope.

### Immunofluorescence (IF) processing

#### anti-p16INK4a+CD31+Stain and p16INK4a+ Stain

Slides were baked slides on their sides for 1 hour at 58-60C in glass staining rack prior to deparaffinization and rehydration. antigen retrieval/ stripping was performed at 90-95C in Citrate buffer AR6 (1:10 type 1 water) (akoya biosciences) for mice for 15 min in vegetable steamer. Slides were rinse slides with Type I water and peroxidase was quenched with fresh 3% H2O2 (Sigma) in H2O for 15 min. Slides were rinsed again with Type I water and blocking buffer was added (10% BSA (Sigma), 0.05% tween 20 (Millipore Sigma) in PBS) for 30 min. Slides were then decanted and incubated with primary antibodies in blocking buffer. For p16INK4a+CD31+ staining this step consisted of 1:1000 Rabbit anti-mouse p16 (abcam) (ab211542) for 30 min in round 1 and 1:2000 Rabbit anti-mouse CD31 (abcame) (ab182981) for 30 min in round 2. Slides were then washed 3x with TBST (Cell Signaling Technology) for 3 min on a shaker. Slides were then incubated for 30 min Rabbit-on-mouse, IgG HPR polymer (RMR622H Biocare Medical) and washed 3x with TBST for 3 min on a shaker. Incubation with Opal diluted in 1x amp diluent (akoya biosciences) then commenced for 10 min. For round 1, 1:150 Opal 570 (akoya biosciences) was added. For round 2 (only if performing p16INK4a+CD31+staining), 1:150 Opal 650 (akoya biosciences) was added. Slides were then washed 3x with TBST 3 min on shaker. If p16INK4a+CD31+ staining, slides then began the process again starting with antigen retrieval/ stripping. After completing all rounds of staining, slides were rinsed with Type 1 water and stained for 5 mins with Spectral DAPI (Akoya Biosciences, FP1490) (2 drops of DAPI per 1ml). A final rinse was than performed followed by application of DAKO mounting medium (Agilent DAKO) and coverslipping (VWR) (1.5 thickness). Slides were allowed to dry overnight covered on bench at 4C.

#### Slide scanning and processing

Stained slides were either scanned using a conventional (Zeiss Apotome microscope) or a spectral microscope (Vectra Polaris Automated Quantitative Pathology Imaging System). Slides imaged on the conventional microscope used the standard Zeiss stitching algorithm. Slides imaged on the spectral microscope were unmixed using inForm Tissue Analysis software with synthetic controls and subsequently stitched using the standard HALO algorithm. All image analysis was performed on the HALO imaging software. Inside and outside annotations for MC38 slides were defined based on a distance of approximately 750um from the surface of the tumor. Cell percentages were calculated using the Indica Labs - HighPlex FL v3.2.1 program.

### Generation of three-dimensional (3D) murine pulmonary maps at cellular resolution

Serial histological images were converted into 3D cellular resolution tissue maps using the CODA platform.^11^ Serial histological images were downsampled from 20x, or approximately 0.5 µm per pixel, to generate 1 and 10 µm per pixel tif images, corresponding to pseudo-10x and pseudo-1x, respectively.

Nonlinear image registration was applied to the pseudo-1x images to generate semi-continuous image stacks from the discrete images. Registration quality was assessed by computing pre- to post-registration tissue image warp (with <20% warp deemed acceptable) and by computing post-registration per-image-pair pixel cross-correlation (with >0.85 deemed acceptable). InterpolAI was used to generate the missing unstained slides to restore microanatomical connectivity in 3D.^37^

Cellular nuclei were detected using Stardist nuclear segmentation applied to the 20x H&E images. For each cell, 20 morphological features were extracted and cell centroids were registered into a volume matrix by applying the transformation matrices calculated on the pseudo-1x H&E images.^38,39^

A semantic segmentation model was trained to recognize nine structures in the murine pulmonary system, consisting of cancer cells, vasculature, bronchioles, nerves, stroma, alveoli, immune clusters, and cartilage.^11,40^ To train the model, a minimum of three images per mouse were randomly selected from the pseudo-10x image stacks. In each image, a minimum of 10 examples of each tissue type were annotated, except in images where certain structures were not found (for example there were no cancer cells in the mock resection lungs). One image from each mouse was used for model testing, with the remainder used for to train a CODA Deeplabv3+ semantic segmentation model. The model was deemed acceptable if the overall accuracy exceeded 90% and the minimum per-class precision and recall exceeded 85%. If the model underperformed, additional annotations were added and the model retrained to improve performance. Following training, the full stacks of pseudo-10x images were segmented, and segmented images were registered using the transformation matrices calculated on the pseudo-1x images to generate a tissue volume matrix.^11,40,41^ Tissues were rendered in 3D, with overlays generated of the cancer cells, bronchioles, vasculature, and immune clusters. Further visualizations were generated using z-projections, where the cumulative voxel density of a given class is calculated down the z-dimension (corresponding to depth in the image stack), colorized, and viewed in two-dimensional format.

To ensure accurate comparison of metastatic burden between mice, all connected clusters in the volumetric matrices labelled as cancer were manually validated by histology review. For each cluster, the bounding box in XYZ was determined, and pseudo-10x resolution image stacks corresponding to this region were exported. Image stacks were loaded into FIJI for review by two different researchers, and each object was either confirmed or noted as a false positive and reassigned as alveoli, stroma, inflammation, or bronchiole based on its appearance. Where researchers agreed, the decision was final, and where researchers disagreed the region was re-reviewed together to come to a consensus decision. Following manual review of cancer clusters, the number and volume distribution of cancer metastases was calculated in each mouse lung. Metastases were differentiated into micro- and macro-metastases using 1.5mm^3^ as a threshold, based on consensus agreement of researchers in histology review.

To identify regions surrounding micro- and macro-metastases, the validated tissue matrix was used to generate 3D distance heatmaps. For each voxel in the volume matrix, the 3D Euclidean distance in µm from vasculature and from cancer metastases was calculated. This distance was overlayed on the pseudo-10x H&E images and saved in .jpg format to enable rapid visualization and review by researchers. These 3D heatmaps enabled automated detection of images containing lung parenchyma near or distant from micro-metastases.

### Integration of serial H&E an Immunohistochemistry (IHC) in 3D

In a subset of mice, multiplex IHC for CD31+CD45+CK19+Vimentin was stained intervening the serial H&E images. IHC images were downsampled from 20x, or approximately 0.5 µm per pixel, to generate 1 and 10 µm per pixel tif images, corresponding to pseudo-10x and pseudo-1x, respectively. CODA nonlinear image registration was performed to integrate the IHC images into the registered H&E pseudo-1x stacks. The pseudo-10x images were aligned by applying the transformation matrices calculated at pseudo-1x, allowing us to integrate the IHC color-intensity with the CODA-labelled tissue types. We extracted the IHC RGB color intensity at regions determined to be cancer cells using the segmentation model in H&E, and compared it to the extracted IHC RGB color intensity at regions determined to be normal lung parenchyma (alveoli). Results were plotted in box-and-whisker plots.

### Integration of serial brightfield and IF images in 3D

In a subset of mice, IF-stained sections were generated intervening the H&E-stained sections. Multiplex IF images were converted to red-green-blue (RGB) color format by assigning P16 to the red channel, CD31 and aSMA to the green channel, and DAPI to the blue channel, and the RGB image was exported at a downsampled resolution of 1 µm per pixel to correspond in scale to the pseudo-1x H&E images. Next, the RGB exported IF images were complemented to better resemble brightfield images where background is white and tissue is darker (rather than IF, where background is black and tissue is lighter). Complementing was necessary for automated nonlinear registration using the CODA platform, as CODA relies on image-to-image pixel cross correlation to determine optimal alignment conditions. These images were registered using the default CODA registration parameters, enabling incorporation of the IF P16, aSMA, CD31, and DAPI channels into the 3D models. ^11^

The registration transforms were applied to the segmented IF masks generated using HALO (see detailed description in the IF processing section above). Registered masks were concatenated into an IF-labelled volume matrix corresponding to the H&E-generated tissue volume matrices and nuclei volume matrices. Using these matrices, the CODA-defined metastases were rendered in 3D and the radial distribution of P16+ senescent cells were calculated surrounding the metastases.

### Slide selection for GeoMx

We selected day 4 to run this analysis, as it was the earliest time point with enough metastases to reliably allow annotations and differential differences from tumor annotations. To include the impact of micro- and large-metastases, we identified micro-metastases using CODA and only considered slides in which they had the largest cross-sectional area. Of these slides, we chose 2 slides from 1 mouse of the tumor bearing and mock control conditions, with the criteria that they contained macro- and micro-metastasis. Of these slides, we narrowed our search down to those containing the maximum cross-sectional area of the micro-metastasis The relevant markers on our panel are aSMA (vasculature), Pan-CK (Epithelial cells), and Syto83 (nuclear). Our goal was to separate out 1) tubular structures less likely to be impacted in the metastatic environment (structures larger than terminal bronchioles, arterioles, and venules) vs 2) parenchymal structures more likely to be impacted by an early metastatic environment (alveoli, immune cells, capillaries, and tumor cells). The parenchymal segment was defined first in the GeoMx collection software as Syto83^+^ Pan-CK^-^ aSMA^-^ cells. The tubular segment was defined second as all remaining Syto83^+^ cells.

Regions of interest (ROIs) were chosen based on the following criteria: 1. Only regions within the parenchyma of the lung were chosen. 2. Annotation areas were chosen to contain both parenchymal and tubular segments. 3. In general, annotations were dispersed between lobes of the lungs. 4. Five tumor annotations were chosen per slide. First, all regions confirmed by CODA to have tumors were annotated. Then, the rest of the 5 annotations were chosen based on regions within a 0.5 mm distance from tumors (in 3D). 5. Six near-distance tumor annotations were chosen per lung (tumor bearing lung). These annotations were chosen based on regions between a 1.5mm-0.5 mm distance from tumors (in 3D). 6. Six far-distance tumor annotations were chosen per lung (tumor bearing lung). These annotations were chosen based on regions between a 1.5mm-2.5 mm distance from tumors (in 3D). 7. Seven mock annotations were chosen per lung (mock resection lung).

### Statistical analysis

#### Tumor growth statistics

plotted as Mean±SEM. Ordinary two-way ANOVA with Šidák multiple comparisons test. * p<0.05, ** p<0.01, *** p<0.001, **** p<0.0001.

#### IF analysis statistics

plotted as Mean±SD. If only comparing two groups, unpaired two-tailed t-test. If comparing three or more groups, ordinary one-way ANOVA with Tukey’s multiple comparisons test. For correlation plots, simple linear regression with 95% confidence band. * p<0.05, ** p<0.01, *** p<0.001, **** p<0.0001.

#### Flow cytometry statistics

plotted as bar graphs with mean ± SD. Mixed-effects analysis with tukey multiple comparisons test. * p<0.05, ** p<0.01, *** p<0.001, **** p<0.0001.

#### GeoMx statistics

QC was run by the Shenderov group. Log2 and padj values were generated using a linear mixed model: ∼ Tags + (1+ Tags | Scan_ID) + (1+ Tags | Scan_ID : Mouse) with Benjamini-Hochberg correction. Tags correspond to each sample, Scan_ID to the slide from which the data was collected, and mouse to the mouse which produced the slides. For near vs far comparison, near vs far annotations were separated in the Tags of the model. For all other analyses these annotations were combined into a tumor adjacent annotation in the Tags of the model. Volcano and GSEA analyses were run on the GeoMx spatial digital profiler (NanoString) platform.

